# Protein structure prediction and design in a biologically-realistic implicit membrane

**DOI:** 10.1101/630715

**Authors:** Rebecca F. Alford, Patrick J. Fleming, Karen G. Fleming, Jeffrey J. Gray

## Abstract

Protein design is a powerful tool for elucidating mechanisms of function and engineering new therapeutics and nanotechnologies. While soluble protein design has advanced, membrane protein design remains challenging due to difficulties in modeling the lipid bilayer. In this work, we developed an implicit approach that captures the anisotropic structure, shape of water-filled pores, and nanoscale dimensions of membranes with different lipid compositions. The model improves performance in computational benchmarks against experimental targets including prediction of protein orientations in the bilayer, ΔΔ*G* calculations, native structure discrimination, and native sequence recovery. When applied to *de novo* protein design, this approach designs sequences with an amino acid distribution near the native amino acid distribution in membrane proteins, overcoming a critical flaw in previous membrane models that were prone to generating leucine-rich designs. Further, the proteins designed in the new membrane model exhibit native-like features including interfacial aromatic side chains, hydrophobic lengths compatible with bilayer thickness, and polar pores. Our method advances high-resolution membrane protein structure prediction and design toward tackling key biological questions and engineering challenges.

**Significance Statement:** Membrane proteins participate in many life processes including transport, signaling, and catalysis. They constitute over 30% of all proteins and are targets for over 60% of pharmaceuticals. Computational design tools for membrane proteins will transform the interrogation of basic science questions such as membrane protein thermodynamics and the pipeline for engineering new therapeutics and nanotechnologies. Existing tools are either too expensive to compute or rely on manual design strategies. In this work, we developed a fast and accurate method for membrane protein design. The tool is available to the public and will accelerate the experimental design pipeline for membrane proteins.

---

Protein design is a powerful tool for elucidating biological mechanisms and engineering new therapeutics. Over the past 20 years, soluble protein design has advanced to atomic level accuracy (1, 2). A remaining challenge is to create tools for membrane proteins (3): a class of molecules that constitute over 30% of all proteins (4) and are targets for 60% of pharmaceuticals (5). There have been several achievements in membrane protein design including a zinc-transporting tetramer Rocker (6), an ion-conducting protein based on the *Escherichia Coli* Wza transporter (7), *β*-barrel pores with increased selectivity (8), receptors with new ligand-binding properties (9, 10), and designed *de novo α*-helical bundles that insert into the membrane (11). For these advances, design of lipid-facing positions often used a native sequence or restricted the chemistry and/or size of amino acids. To apply membrane protein design to addressing biological questions and engineering challenges, tools must sample a realistic distribution of amino acids tied with the diverse lipid composition.

The foundation of computational design tools is the energy function: a mathematical model of the physical rules that distinguish native from non-native membrane protein conformations and sequences. Currently, most computational studies of membrane proteins are molecular dynamics simulations with an all-atom lipid bilayer. In this conception, the lipid molecules are represented explicitly using force fields such as AMBER(12), CHARMM (13, 14), Slipids (15), or GRO-MOS (16), and the protein-lipid interactions are scored with a molecular mechanics energy function. All-atom models are attractive because they can feature hundreds of lipid types toward approximating the composition of biological membranes (17). With current technology, detailed all-atom models can be used to explore membrane dynamics for hundreds of nanoseconds (18): the time scale required to achieve equilibrated properties on a bilayer with approximately 250 lipids (19). Coarse-grained representations such as MARTINI (20), ELBA (21), and SIRAH (22) reduce computation time by mapping atoms onto representative beads. As a result, simulations have explored dynamics up to the millisecond time scale to access features of membrane organization and large protein domain motions (23).

Implicit solvent models enable simulations to reach longer timescales required to investigate biologically-relevant conformational and sequence changes. Instead of using explicit molecules, implicit methods represent the solvent as a continuous medium (24, 25), resulting in a 50—100-fold sampling speedup (26). The most detailed implicit model is the Poisson-Boltzmann (PB) equation, which relates electrostatic potential to dielectric properties of the solvent and solute through a second-order partial differential equation (27). Numerical solvers have enabled PB calculations on biomolecular systems (28, 29); however, these calculations do not scale well. To reduce computational cost, the Generalized Born (GB) approximation of the PB equation treats atoms as charged spheres (30). GB methods represent the low-dielectric membrane through various treatments ranging from a simple switching function (31) to heterogeneous dielectric approaches (32, 33). However, evaluating the GB formalism is still computationally expensive.

A popular approach to overcoming the computational cost of solvent electrostatics models is the Lazaridis implicit membrane model (IMM1; (34, 35)): a Gaussian solvent-exclusion model that uses experimentally measured transfer energies of side-chain analogues in organic solvents to emulate amino acid preferences in the bilayer (36). IMM1 has been applied to various biomolecular modeling problems including studies of antimicrobial peptides (37), *de novo* folding (38, 39), and *de novo* design of transmembrane helical bundles (11). However, organic solvent slabs differ from phospholipid bilayers because lipids are thermodynamically constrained to a bilayer configuration, resulting in a unique polarity gradient that influences side chain preferences (40–42). A possible alternative is to directly calculate amino acid preferences by deriving statistical potentials from a database of known membrane protein structures (38, 43, 44). Yet, statistical potentials are also limited by the scarcity of membrane protein structures and do not capture the physiochemical properties of the membrane.

In this work, we developed, implemented, and tested a biologically-realistic energy function for membrane protein structure prediction and design. We first derived and validated the energy function parameters from experimental and computational modeling of phospholipid bilayers to capture biologically-important membrane features. Next, we tested the model on four benchmarks: (1) prediction of protein orientations in the membrane, (2) ΔΔ*G* of mutation calculations, (3) native structure discrimination and (4) native sequence recovery. We investigated properties of the *in silico* designed membrane proteins including the amino acid composition. Finally, we share several design anecdotes that exhibit native-like membrane protein features including interfacial aromatic side chains, hydrophobic lengths compatible with different lipid compositions, and polar pores.

## Results

### Biologically-realistic implicit membrane model

We developed a biologically-realistic implicit membrane model inspired by Lazaridis’ implicit model (IMM1; (35)). Similar to IMM1, the membrane is modeled as a continuum of three phases: an isotropic phase representing bulk lipids, an isotropic phase representing bulk water, and an anisotropic phase representing the interfacial region. To accurately model the polarity gradient and dimensions of native membranes, we derived new equations and parameters from biophysical measurements. The result is a new energy term called Δ*G*_memb_ that computes protein stability given the water-to-bilayer transfer energy 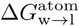 of atomic groups *a* and the fractional hydration *f*_hyd_:

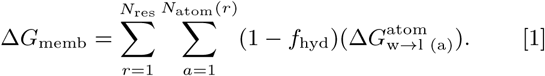

The parameter 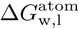 captures the thermodynamics of protein-lipid interactions. We derived 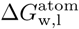 from the Moon & Fleming side-chain hydrophobicity scale (45) because the energies were measured in bilayers with phospholipids, a major component of biological membranes (46). Following Lazaridis’ formalism (47), the function *f*_hyd_ captures the three-dimensional shape of the implicit membrane as a dimensionless number that describes the phase given the position of an atomic group. When an atomic group is exposed to the lipid phase, *f*_hyd_ = 0; whereas when an atomic group is exposed to the water phase, *f*_hyd_ = 1.0. The transition between the two isotropic phases is modeled by a composition of functions:

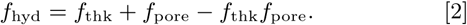

The function *f*_thk_ (Eq. 3, see Methods) models the transition between the water and lipid phase along the *z*-axis, and is thus an implicit representation of the hydrophobic thickness. We developed parameters for *f*_thk_ by fitting to molecular dynamics simulations and scattering density profiles of phospholipid bilayers. The result is a logistic curve that depends on two parameters. We derived parameters for fourteen phospholipid bilayer compositions (Table S4-5). The membrane thickness can be derived by setting *f*_thk_ = 0.5 (Fig. 1A-B).

**Fig. 1.**
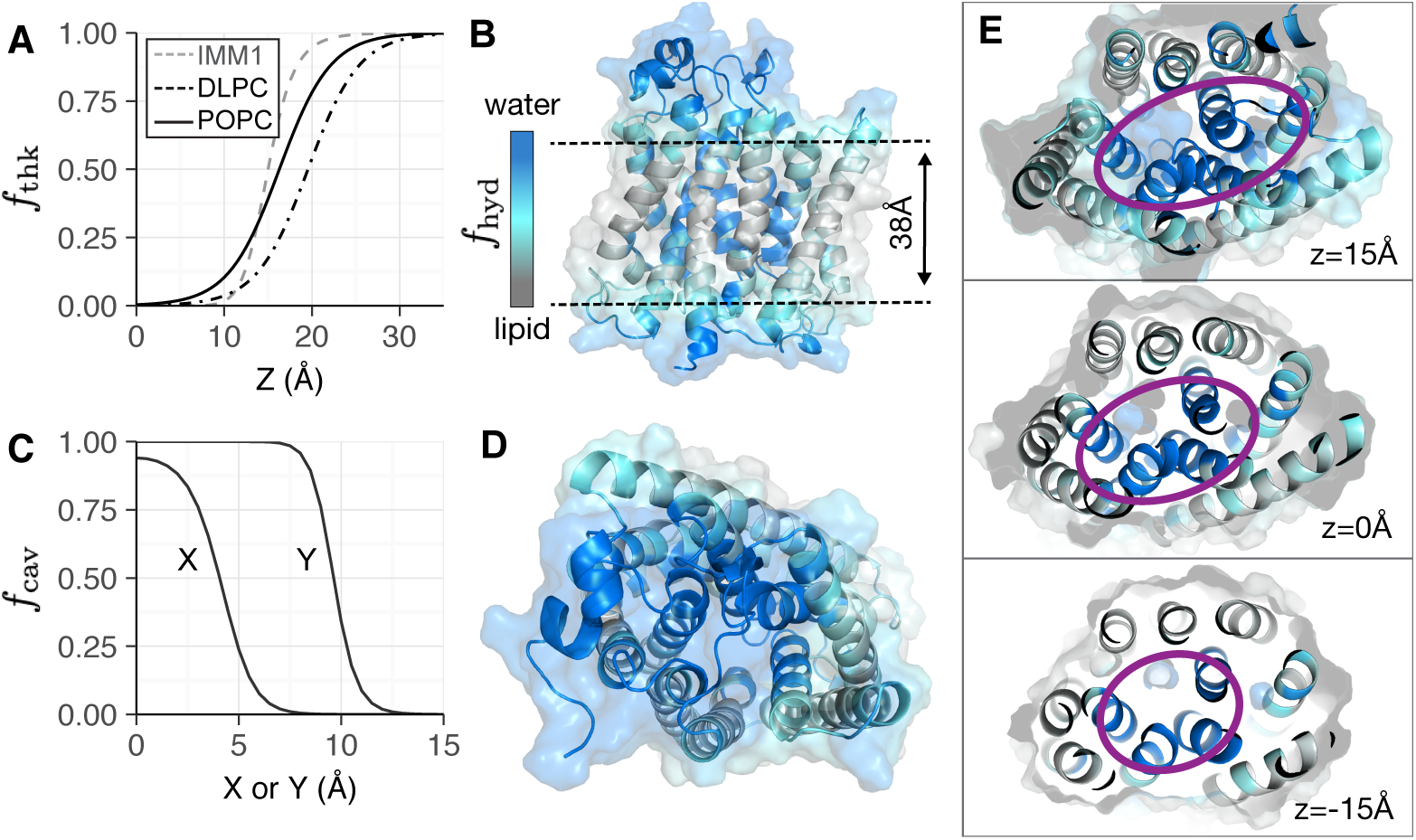
The implicit membrane is modeled as three phases: two isotropic phases for water and lipid and a transition region that represents the interfacial head groups. (A) The transition between phases in the *z*-dimension is modeled by a logistic curve which can be parameterized for different lipid compositions. Example curves for DLPC (dot-dash, black) and DOPC (solid, black) are shown in comparison to the sigmoid curve used in IMM1 (dashed, gray). (B) Implicit solvent phases for the Ammonium transporter Amt-1 (PDB 2b2f) in the *z*-dimension. The water phase is shown in blue, the interface is in teal, and the lipid is in gray. (C) The transition between phases due to an elliptical pore is modeled by a sigmoid curve. (D) Top view of implicit solvent phases due to a pore in Amt-1 with the same coloring scheme as B. The three panels of (E) demonstrate the variation in pore shape (purple) for different *x - y* cross sections along the *z*-axis.

The function *f*_pore_ defines the shape of a water-exposed pocket or channel (Fig. 1C-E). Previously, Lazaridis developed a cylindrical model of pores for *β*-barrel proteins (47). This geometric assumption is straightforward for *β*-barrel proteins; however, *α*-helical protein pores require varied geometric descriptors such as cones, cylinders, and ellipses (48). To accommodate, we created a model that approximates pores as elliptical tube with varying cross sections. This parameterization allows the model to describe cavities that do not penetrate through the membrane and pores that constrict, expand, or twist relative to *z*. The energy function accounts for the pore by first calculating a relative radius, *g* _radius_ (Eq. 5). The transition between the two phases is modeled by a sigmoid curve *f*_pore_ (Eq. 6) with two parameters: *g*_radius_ and the transition steepness *n* (default *n* = 10).

We integrated our model into the current all-atom energy function for modeling soluble proteins in Rosetta, called REF15 (49). REF15 computes macromolecular energies through a linear combination of terms for van der Waals, solvation, electrostatics, hydrogen bonding, backbone- and side-chain interactions. To account for the membrane environment, we added Δ*G*_memb_ with an empirically determined weight of 0.5. The resulting energy function, called *franklin2019*, is given by Δ*E*_franklin2019_ = Δ*E*_REF15_ + Δ*G*_memb_.

### Computational benchmark performance of the biologically-realistic implicit membrane

We evaluated the accuracy of *franklin2019* using four computational benchmark tests against experimental targets. The tests were designed to evaluate an energy function’s ability to replicate measured membrane protein stabilities and perform accurate structure prediction and design. We compared the performance of *franklin2019* to three existing models: (1) an implicit membrane parameterized from the behavior of side-chain analogues in organic solvents (**M07**; (39)), (2) a knowledge-based model that captures depth-dependent amino acid preferences (**M12**; (50)), and (3) the Rosetta all-atom energy function for soluble proteins (**R15**; (49, 51)). For brevity, we will refer to *franklin2019* as **M19**. We chose these models because the low computational cost enabled evaluation with structure prediction and design tests.

#### Test #1: Partitioning of transmembrane peptides into the implicit membrane

The water-to-bilayer transfer energy of a protein Δ*G*_w,l_ thermodynamically stabilizes the protein in the membrane. Therefore, implicit membrane energy functions must accurately estimate the value of Δ*G*_w,l_. We investigated the partitioning properties of **M19** by computing the water-to-bilayer partitioning energies and full energy landscape of trans-membrane peptides as a function of membrane orientation. We define orientation by two degrees of freedom: helix tilt angle and membrane depth (Fig. 2A).

**Fig. 2.**
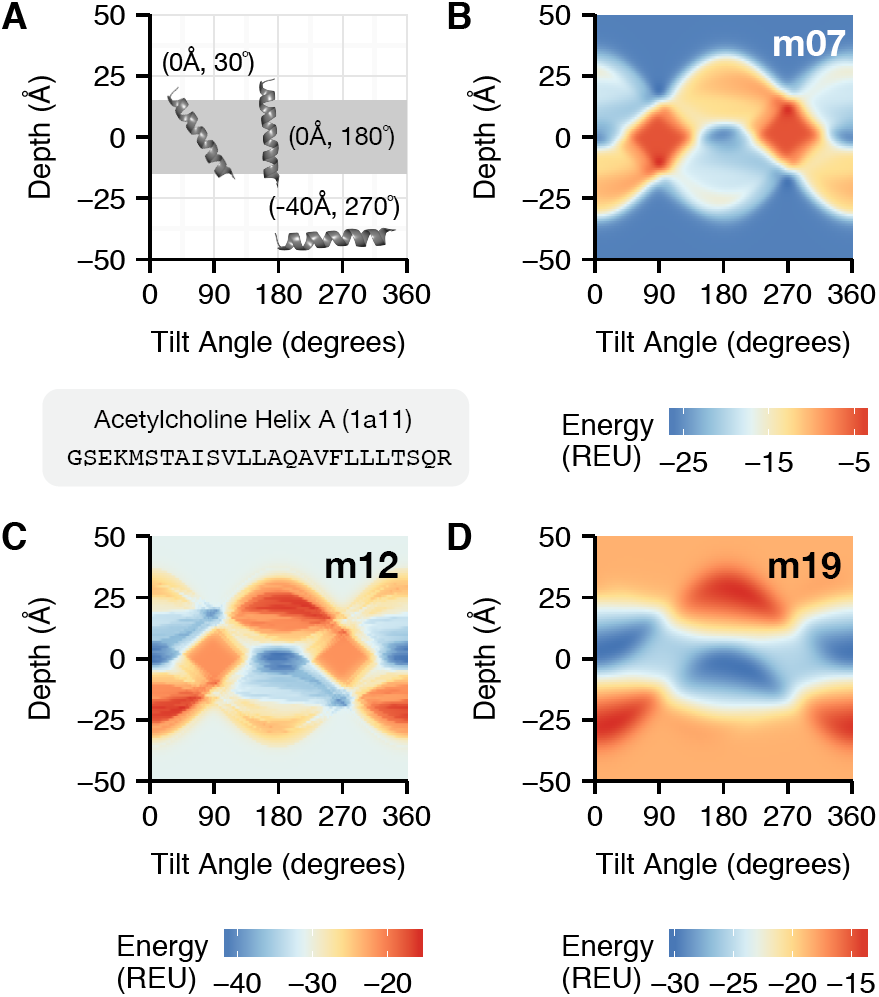
Orientation-dependent energy landscape of the acetylcholine M2 helix (PDB 1a11). The orientation-dependent landscape represents the relative energies of a protein at all possible tilt angles and depths. (A) Sample peptide orientations in the energy landscape grid. (B, C, and D) are energy landscape plots with the energy colored from blue (favorable), to yellow (moderate), to red (unfavorable) computed by the M07, M12, and M19 energy functions, respectively.

Remarkably, **M19** was the only energy function for all five peptides in the membrane to predict a favorable Δ*G*_w,l_ (Table 1, see Methods). To investigate further, we mapped all possible orientations to their **M07, M12**, and **M19** energies. An example for the acetylcholine M2 peptide (1a11) is shown in Fig. 2. The **M07** energy landscape (Fig. 2B) has three small, low energy wells and they are isoenergetic with the water phase. This behavior is nonphysical and is likely an artifact of inconsistent parameterization. In contrast, the lipid phase is more thermodynamically favorable than the water phase for both **M12** (Fig. 2C) and **M19** (Fig. 2D). Ultimately, **M19** is the most native-like because the landscape is smooth and continuous: a requirement for high-resolution all-atom structural modeling.

**Table 1.** **Partitioning energy from water to lipid**, Δ*G*_**w,l**_

| Target (PDB ID) | $\Delta G_{w,l}^{M07}$ (REU) | $\Delta G_{w,l}^{M12}$ (REU) | $\Delta G_{w,l}^{M19}$ (REU) |
| --- | --- | --- | --- |
| 1a11 | 1.32 | -10.41 | -11.44 |
| 1mp6 | 0.26 | -20.46 | -15.77 |
| 1pje | -4.46 | -25.42 | -15.24 |
| 2nr1 | 14.41 | 7.55 | -7.47 |
| WALP23 | -7.23 | -29.41 | -17.80 |

The energy landscapes for the remaining four targets are given in Fig. S5. We found that the M2 proton channel segment (Fig. S5A) and NMDA glutamate receptor segment (Fig. S5B) demonstrate similar characteristics to acetylcholine. Namely, that **M19** generates the smoothest landscape with low energy wells at realistic peptide orientations. Here, a reasonable orientation is when the peptide vertically spans the transmembrane region. Interestingly, the **M07** energy landscape features were different for the VPU-forming domain (Fig. S5C) and WALP23 (Fig. S5D) targets. While the landscapes remain rugged, there is an unfavorable energy gradient from the aqueous to the lipid phases. It is unsurprising that WALP23 is a compatible peptide with **M07** because it is the only non-naturally occurring peptide in the group. Rather, the sequence was engineered to favorably partition into a wide variety of membrane mimetics (52). While it is less clear why VPU-forming domain is compatible, likely the shorter peptide length reduces the barrier to inserting charged termini.

We also evaluated the ability of **M19** to discriminate native and non-native orientations in the membrane. Initially, we compared experimentally measured tilt angles (53) with calculated tilt angles corresponding to energy minima (see Methods). However, no energy function could reproduce tilt angles within five degrees. We hypothesized this is because we do not evaluation the entropic contribution to peptide tilt (54). Instead, we asked if the minimum energy orientation is native like: the peptide tilt angle is under 45°and its membrane depth is ±5 Åfrom the center, as supported by solid-state NMR studies that demonstrate native peptides are slightly tilted relative to the membrane normal while still spanning the membrane (55). **M19** found a reasonable orientation for four of five targets (Table S9). The **M12** identified a reasonable orientation for three of five peptides and **M07** only identified a reasonable orientation for WALP23. Thus, **M19** demonstrated the best performance.

#### Test #2: Predicting the ΔΔ*G* of mutation

Predicting changes in protein stability upon single amino acid substitutions at lipid exposed positions informs predictions of the effects of genetic mutations and *de novo* protein design. We evaluated the ability of **M19** to capture the change in protein stability upon mutation, called ΔΔ*G*^mut^, by comparing experimentally measured values with computational predictions. Here, we used a dataset of mutations at position 111 on outer membrane palmitoyl transferase (PagP) (56)). The dataset contains mutations from the host amino acid to all 19 other canonical amino acids. A summary of prediction accuracy relative to the experimentally measured values is given in Fig. 3. The raw predicted values are also listed in Table S10 for PagP. Calculated energies are given in Rosetta Energy Units (REU).

**Fig. 3.**
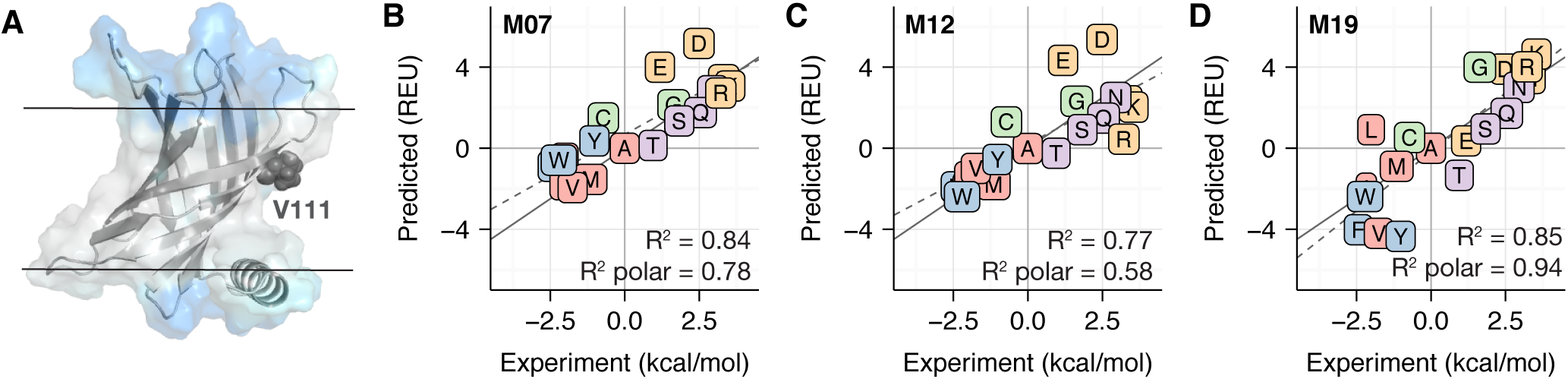
Comparison between computationally predicted and experimentally measured ΔΔ*G*^mut^ values. For all correlation plots (B-D), proline is not shown due to steric clashes resulting in a large ΔΔ*G*^mut^ value. The dotted gray line is the line of best fit and the solid gray line is *y* = *x*. In addition, amino acids are colored according to the following categories: charged (orange), nonpolar (red), aromatic (blue), polar (purple), special case (green). (A) Structure of the PagP scaffold (PDB 3gp6) with the mutation site V111 highlighted in dark grey. The implicit solvent phases in A are colored in a similar manner to Fig 1. The ΔΔ*G*^mut^ predictions for mutations in PagP by M07, M12, and M19 are shown in panels B, C, and D respectively.

The correlation between **M19** predicted and experimentally measured ΔΔ*G*^mut^ values was R^2^ = 0.85. Note, the ΔΔ*G*^mut^ for proline was excluded for all three energy functions because steric clashes resulted in large values. While prediction accuracy was improved relative to **M12** (R^2^ = 0.77), accuracy was comparable to **M07** (R^2^ = 0.84). We were surprised that **M07** and **M19** demonstrated similar predictive ability. This is because a second set of measurements in OmpLA (45) correlates well with PagP measurements but not with **M07** predictions (57). According to Marx *et al.*, the largest deviations were for side chains containing polar atoms. We therefore recalculated the correlation coefficient for polar and charged side chains. Here, the correlations were 0.78, 0.58, and 0.94 for **M07, M12**, and **M19** respectively. We were encouraged by these results because they demonstrate the ability of our scale to capture the behavior of polar side chains within the bilayer.

To dive deeper, we examined the incorrect predictions, defined as cases where the calculated ΔΔ*G*^mut^ deviates by more than 1.5 REU from the measured value. For **M19**, we found that G, T, V, and L were inaccurate. From the component energies (Tables S12-14), we learned that the predictions are incorrect due to a large Lazaridis-Karplus solvation energy and our water-to-bilayer energy. In the future, we anticipate that improved experimental techniques will provide more data to re-balance these energy terms.

#### Test #3: Discrimination of native structures from decoys

Identification of native-like structures in an ensemble of candidate structures is a key function of biomolecular modeling energy functions. We evaluated native structure discrimination on molecular-dynamics-based ensembles of five targets: bacteriorhodopsin (Brd7), fumarate reductase (Fmr5), lactose permease (LtpA), rhodopsin (RhoD), and V-ATPase (Vatp) (58). The root-mean-squared-deviation (RMSD) between the native and the candidate models ranged between 4-15Å. First, we compared the scores of refined native models and decoys. We found that for all targets except LTPA, all of the energy functions distinguished the native structure from the decoys. Qualitatively, we observed that R15 and M19 decoys formed deeper energy funnels with near-native decoys at lower energies than distant decoys. To quantitatively evaluate decoy discrimination, we computed the *W*_RMS_ for all targets (Table 2). In addition, a mapping of energy vs. RMSD for each target refined by each energy function is given in **Fig. S8**.

**Table 2.** Weighted RMSD of refined and rescored candidate models by each energy function

| Target | R15 | M07 | M12 | M19 |
| --- | --- | --- | --- | --- |
| Brd7 | 4.76 | 4.94 | 6.38 | 5.15 |
| Fmr5 | 5.31 | 5.21 | 9.78 | 4.98 |
| LtpA | 4.53 | 4.47 | 4.69 | 4.62 |
| RhoD | 4.38 | 4.16 | 4.10 | 4.43 |
| Vatp | 4.30 | 4.37 | 4.22 | 4.35 |
| Average | 4.66 | 4.61 | 5.83 | 4.70 |

The *W*_RMS_ data confirmed poor discrimination by **M12** with an average *W*_RMS_ of 5.83Å. This is consistent with the funnel plots in Fig. S8 where M12 does not assign low energies to near-native decoys. Interestingly, the remaining energy functions demonstrate similar performance, with *W*_RMS_ values differ by less than 1Å. The average *W*_RMS_ values are also within 1Åof the lowest RMS decoy. Thus, **M07, M19**, and **R15** can reliably distinguish near-native from non-native decoys.

We were surprised that the new implicit membrane model did not have a larger impact on native structure discrimination. For this, we provide two hypotheses. First, there may be insufficient near-native decoys to fully capture the energy landscape. This is likely because there were less than 130 decoys for each target, whereas 1,000-5,000 are required for sufficient conformational sampling. A second hypothesis is that the non-membrane energy terms drive high-resolution decoy discrimination.

#### Test #4: Native sequence recovery

A fourth test evaluates sequence recovery: the fraction of amino acids recovered after performing complete redesign on naturally occurring proteins. High sequence recovery has long been correlated with strong energy function performance for soluble proteins (59). We therefore repeated this test in the context of our membrane protein energy function. We computed low free energy sequences for a test set of 133 *α*-helical and *β*-barrel membrane proteins. Overall, 31.8% of the amino acids designed by **M19** were identical to the native amino acid (Fig. 4A). The soluble protein energy function **R15** recovered the second highest percentage of amino acid positions at 29.9%. In contrast, the two existing implicit membrane models lagged behind with **M07** at 26.5% and **M12** at 26.7% (Fig. 4A). The individual amino acid recovery rates were also revealing. Here, **M19** and **R15** recovered all 20 amino acids at rates above random; whereas **M12** recovered 19 and **M07** only recovered 14.

**Fig. 4.**
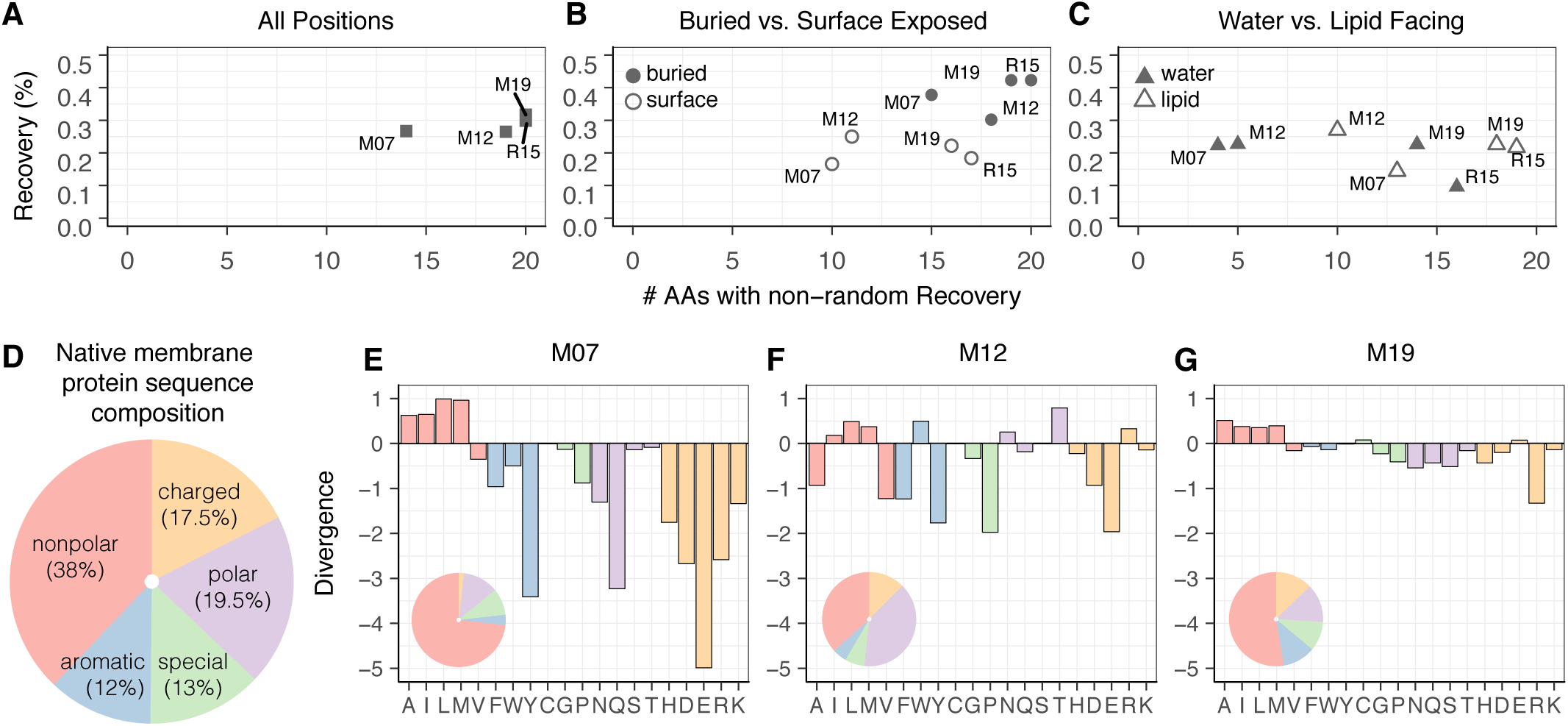
Properties of membrane protein sequences designed by R15, M07, M12, and M19 relative to their native counterparts. Sequence recovery vs. *N*_AA_ for all positions (A), buried vs. surface-exposed positions (B), and water vs. lipid exposed positions (C). (D) Amino acid composition of the native sequences in the benchmark set. Divergence of the amino acid distribution in proteins designed by M07 (E), M12 (F), and M19 (G) relative to the amino acid distribution in native membrane proteins. A positive value indicates that an amino acid is over-enriched, whereas a negative value indicates that an amino acid is under-enriched. Values are given on a logarithmic scale. An amino acid composition pie chart for sequence designed by each candidate energy function is also shown in the bottom left hand corner of the divergence plots.

To examine the influence of different solvent environments, we recomputed recovery over subsets of residues. First, we compared buried vs. solvent-exposed side chains (Fig. 4B). For all energy functions, recovery was significantly higher for buried side chains than solvent-exposed side chains, as noted in previous studies due to higher packing density (59). On the surface, **M12** recovered 25% of acid positions, slightly higher than the 22% recovery rate by **M19**. However, **M19** recovered 16 amino acids at rates above random; whereas, **M12** recovered only 12 amino acids. In essence, **M19** gets the overall answer correct slightly less frequently; however, it is better at getting more amino acid types correct.

Next, we examined sequence recovery differences between side chains facing the water and lipid phases (Fig. 4C). In the lipid phase, all membrane energy functions recovered nearly the same fraction of amino acids. The main differentiating feature is the number of amino acids recovered with greater than random probability. Whereas **M07** and **M12** recovered four and five amino acids respectively,**M19** recovered 14 amino acids. We observed a similar trend in the water phase. Here, **M12** has the highest overall sequence recovery rate of 27%, next to **M19** with a recovery rate of 23%. However, M12 only recovers 10 amino acid types whereas M19 recovers 14. These results reveal that early energy functions used a rudimentary design strategy: prioritizing only some amino acid types. In contrast, **M19** is capable of designing more chemically diverse sequences.

### Designed membrane proteins exhibit native-like features

The sequence recovery experiment enables us to study properties of *in silico* designed membrane proteins. These properties are crucial for demonstrating that the implicit model has native membrane properties and is capable of facilitating realistic design experiments. Below, we examine various sequence and structural features important for membrane protein stability and function.

#### Amino acid distribution in designed proteins mirrors the native distribution

We examined the distribution of amino acids in design protein sequences relative to their native counterparts. Specifically, we measured the Kullback-Leilber (*D*_KL_) divergence (Eq. 9, see Methods) on our membrane protein dataset. A negative *D*_KL_ value indicates that sequences are under-enriched in specific amino acid types; whereas, a positive *D*_KL_ value indicates that sequences are over-enriched. An ideal KL value is zero. Remarkably, sequences designed by **M19** are near-native with *D*_KL_ = −2.7. This is in stark contrast to sequences designed by **M07** and **M12** which are strongly divergent from native membrane protein sequences, with *D*_KL_ = −24.6 and *D*_KL_ = −26.6 respectively.

To learn more about the design implications of each energy function, we computed the KL for each amino acid type (Fig. 4D-G) and compared to the composition of amino acids in the native set. The **M07** sequences are over-enriched in non-polar amino acids and under-enriched in all other categories. The deficits are large with under-enrichment values ranging from 10^−2^ to 10^−4^. The **M12** sequences are less skewed with the magnitude of under-enrichment deficits ranging between 10^−1^ and 10^2^. However, there is still a large over-enrichment of non-polar amino acids including I, L, and M, as well as W and T. In contrast, the distribution of amino acids in **M19** sequences is comparable to the native distribution, with the magnitude of under- and over-enrichment values ranging between 10^1^ and 10^−1^. Thus, **M07** and **M12** employ a rudimentary design strategy: only choosing non-polar amino acids guaranteed to be compatible with the greasy membrane environment. The **M19** model does not rely on this assumption and can design every amino acid type within each phase. As a result, **M19** designs proteins with an amino acid distribution that is close to the native membrane protein sequence composition. We thus expect that **M19** will more accurately evaluate the effects of genetic mutations on protein stability. Further, the diversified sequences will enable designed membrane proteins to achieve a broader range of architectures and functions.

#### Three-dimensional membrane geometry enables design of polar pores

We were interested to see whether a three-dimensional implicit membrane shape facilitates accurate protein design. To do so, we investigated the native and designed sequence of the scaffold protein voltage-dependent anion channel 1 (VDAC1; PDB 3emn; Fig. 5). The native sequence of this *β*-barrel protein pore is rich in charged amino acids. In the two-dimensional membranes used by **M07** and **M12**, the pore-facing residues are designed as if they are in the lipid phase; and as a result, the designed sequences are rich in non-polar amino acids. In contrast, the three-dimensional implicit membrane geometry treats pore-facing residues as exposed to the water phase; thus, the designed sequence contains both polar and charged amino acids. These positive features are reflected in the sequence for this specific target. Here, **M19** exhibits the highest recovery over all surface residues, lipid-facing, and aqueous-facing residues when compared with other energy functions. This results suggests the potential of **M19** to perform accurate design on both the lipid-facing and water-filled-pore facing surfaces.

**Fig. 5.**
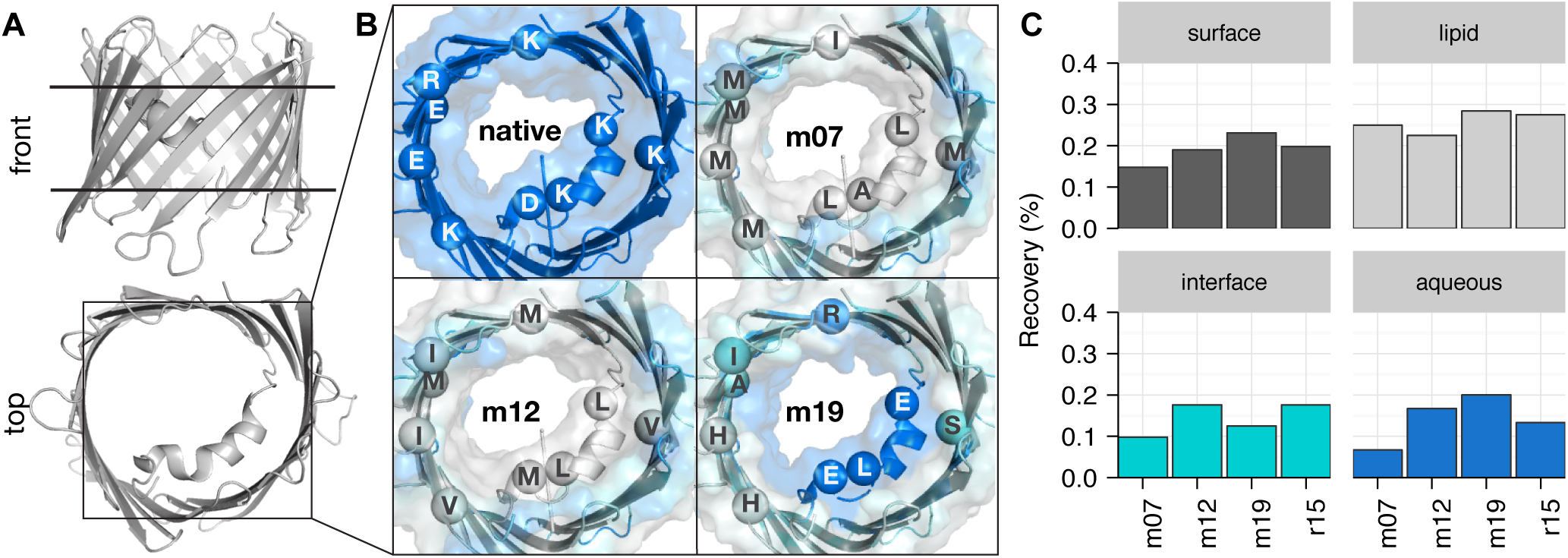
An *in silico* redesigned *β*-barrel membrane protein with a polar aqueous pore. (A) Structure of the design scaffold protein voltage-dependent anion channel VDAC1 (PDB 3emn) from a lateral and top view. (B) Sequence composition and solvation properties of the pore redesigned by the m07 (top right), m12 (bottom left), and m19 (bottom right) energy functions in contrast to the native sequence (top left). The m07 and m12 treat the pore as lipid-exposed resulting in a non polar sequence. In contrast, the m19 energy function calculates a custom pore shape resulting in a polar pore sequence. (C) Recovery of the native 3EMN sequence upon redesign. In contrast to other energy functions, the m19 recovers the most native sequence for the total surface, lipid-exposed, and aqueous-solvated residues.

#### Biologically-relevant lipid composition parameters improve per-target sequence recovery

Finally, we were eager to explore whether implicit membrane parameters for different lipid compositions can improve design outcomes. This question is difficult to evaluate because the host membrane composition of proteins is not always known. At the same time, this question is crucial because of the long-standing criticism that implicit membrane models do not accurately capture the properties of different lipid membrane compositions. In this work, we investigated this question anecdotally by examining two examples from our membrane protein design dataset.

First, we examined the *β*-barrel protein scaffold outer membrane transporter FecA from *Escherichia coli*. The outer membranes of gram-negative bacteria are significantly thinner than eukaryotic plasma membranes. We therefore hypothesized that sequence recovery of lipid-facing residues in this protein would be higher in a thinner membrane. To test this hypothesis, we again searched for low energy sequences in an **M19** membrane with either DLPC or POPC parameters. Encouragingly, the recovery of lipid facing residues in this protein was 33% in DLPC in contrast to 28% in POPC. We also repeated this test on the *α*-helical protein scaffold VCX1 calcium/proton exchanger from *Saccharomyces cerevisiae*. Here, we expected the reverse trend: improved design in a POPC membrane over DLPC. Again, the design results followed: 22% sequence recovery in DLPC and 29% in POPC. These results demonstrate that lipid composition parameters facilitate more biologically-realistic structure prediction and design.

In addition, there was an inevitable question that we wanted to ask about our *β*-barrel protein scaffold. Experimental studies have long demonstrated that *β*-barrel membrane proteins have high concentrations of aromatic side chains near the interfacial head groups (60). While the thermodynamics of this phenomena are not completely understood, it is suggested that stacking of the aromatics nearby polar head-groups stabilizes the protein (61). Thus, we asked the questions: does **M19** also design aromatics near the anisotropic phase representing interfacial head groups? To answer, we calculated the apparent membrane thickness according to the average positions of aromatic side chains in native and designed FecA (Fig. 6). We found that **M19** designed with a larger apparent thickness in POPC rather than DLPC membranes. Notably, the DLPC aromatic thickness is near the native aromatic thickness. While still anecdotal, these results suggest that **M19** designs proteins with native-like features.

**Fig. 6.**
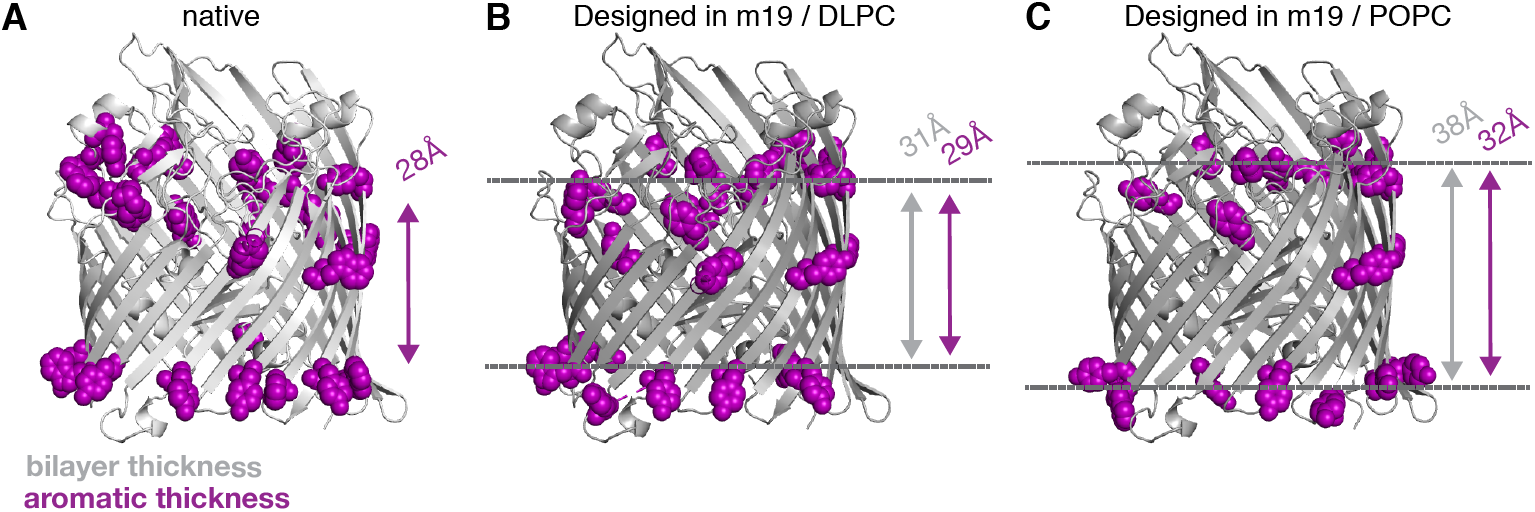
*In silico* redesigned *α*-helical and *β*-barrel membrane proteins in native-like lipid compositions. (C) Structure of the design scaffold Outer Membrane Transporter FecA (PDB 1kmo) from *Escherichia coli* redesigned with the M19 energy function with DLPC and POPC parameters. The native scaffold is colored in light grey and the design scaffolds are colored in dark grey. Aromatic amino acids near the interface (>7 and < 25 from the center) are colored in light pink. The grey arrow shows the bilayer thickness and the pink arrow shows the thickness according to the average position of interfacial aromatic residues. The thickness from the DLPC design best matches the native and in (D) results in the highest recovery of lipid facing residues in the transmembrane region. DLPC is closest to the thickness in *E. coli*.

## Discussion

In this work, we developed, implemented, and tested a biologically-realistic energy function for membrane protein structure prediction and design. The energy function, called *franklin2019*, uses an implicit approach to represent the anisotropic structure, water-filled pore shape, and nanoscale dimensions of membranes with varied lipid composition. Through computational benchmarking, we demonstrated that the model can replicate experimentally measured protein stabilities and accurately perform biomolecular modeling tasks. *Franklin2019* outperforms both the Rosetta soluble protein and membrane protein energy functions.

In addition, we investigated the properties of native and designed membrane protein sequences. Remarkably, the distribution of amino acids in sequences designed with *franklin2019* is comparable to the amino acid distribution in native membrane protein sequences. This result is in stark contrast to sequences designed by prior membrane energy functions that are over-enriched in non-polar amino acids. Proteins designed by *franklin2019* exhibit other native-like features including polar pores, aromatic amino acids near interfacial head groups, and hydrophobic match with specific lipid compositions. Together, these features demonstrate the potential of *franklin2019* to advance high-resolution membrane protein structure prediction and design.

Our goal was to develop an implicit model that captures features of biological membranes. We built upon several prior studies that improved the realism of the hydrophobic slab to make it behave like a native bilayer. For instance, Lazaridis added a cylindrical pore (47), parameters for anionic lipid compositions (62), and corrections for charged and buried groups (63). Panahi & Feig developed a generalized born membrane model with adjustable thickness (33). An important differentiating feature of our work is that our model was developed using data from native proteins in native bilayers, thus removing a layer of assumptions and closes gaps in the design cycle of experiment and computation.

In the future, more detail may improve the realism of implicit models. For instance, the lipid composition of the outer membrane of gram-negative bacteria is asymmetric (64). The membrane also bends and curves to accommodate the hydrophobic surface of proteins (65). A further challenge is accounting for local properties such as specific protein interactions with lipids and cholesterol, which may be captured by a hybrid implicit-explicit approach such as SPadES (66) or HMMM (67). An open question is how to account for mechanical properties such as lateral pressure and strain due to local curvature. In these scenarios, it is most likely that implicit membrane simulations will compliment information from emerging membrane protein modeling tools and MD simulations to investigate structure, dynamics, and function.

We evaluated our implicit membrane model using sparse, high-resolution experimental data. This approach contrasts soluble protein energy function evaluation, where there is an abundance of thermodynamic and spectroscopic measurements of small molecules (68–70) and high-resolution protein structures (71). To overcome the possibility of over-fitting, we limited the validation data to high-quality measurements. For instance, we did not use crystal structures ≥ 3 Å resolution or ΔΔ*G*_mut_ values that were not measured in a reversible system. Further, we benchmarked our energy function against both thermodynamic and structure prediction data. Previous studies have evaluated membrane energy functions on a single test such as tilt angles (53), native structure discrimination (58, 72), predicting hydrophobic lengths (73), ΔΔ*G* prediction (74) and sequence recovery (75). Simultaneously performing the benchmarks enables a well-rounded evaluation of the energy function for diverse biomolecular modeling tasks. As more high-quality data emerges, we envision the application of more robust fitting techniques including machine learning and optimization (51, 76).

Through the goal of developing a new energy function, our study interrogated fundamental questions about the design rules for native membrane proteins. First, the transfer energies are derived from a thermodynamic hydrophobicity scale measured in phospholipids. The high sequence recovery rate demonstrates the importance of thermostability and bulk phospholipid chemistry in constraining membrane protein sequences. Further, previous work relied on narrow membrane protein design rules such as enrichment of leucine side-chains in the hydrophobic core. We demonstrated that native membrane protein sequences are diverse and not constrained to hydrophobic amino acids. As a result, the energy function uses the full palette of amino acid chemistries during design.

In summary, we developed a biologically-realistic energy function for membrane protein structure prediction and design. The energy function is implemented within the Rosetta software and can be used for a wide range of macromolecular modeling tools. By pursuing a balance of efficiency and accuracy, we anticipate that the implicit membrane will enable high-throughput and high-resolution membrane protein structure prediction and design. Importantly, this model transforms once protein-centric tools to techniques that can predict and design structures tied with varied biological lipid compositions.

## Materials and Methods

### Development of the implicit membrane model

#### Derivation of 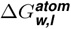 values

The Moon & Fleming hydrophobicity scale provides a set of water-to-bilayer transfer energies 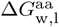 for the 20 canonical amino acids (45) measured in the reversibly folding OmpLA scaffold. We used regression to derive energies that correspond to atom types (Table S1), called 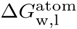. Specifically, least-squares fitting was applied solve the equation **Ax** = **b**; where, **A** is a matrix of atom type stoichiometric coefficients (Table S2), **b** is the vector of 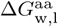 values, and **x** is the desired vector of 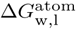 values. Matrix rows for glycine, alanine, and proline were excluded to avoid over fitting. The resulting 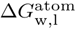 values are in Table S3.

#### Molecular dynamics simulations of phospholipid bilayers

All-atom molecular dynamics simulations were performed to extract properties of membranes with different phospholipid compositions. We simulated phospholipid bilayers with hydrocarbon tails between 12-18 carbons long and either a phosphatidylethanolamine (PE), phosphatidylcholine (PC) or phosphatidylglycerol (PG) head group (Table S4). The exceptions were DPPC and DMPG because the liquid-to-gel phase transition temperatures are above physiological temperature (77, 78). CHARMM-GUI (79) was used to configure each bilayer system with 75 lipids in each leaflet, 22.5Å of water on each side, and 0.1 M NaCl. Simulations were performed using the NAMD molecular dynamics engine (80) at a constant pressure of 1 atm and a temperature of 37°C. We used the CHARMM36 (13) force field for lipid and the TIP3 model for water. The simulations were equilibrated with restraints according to the procedure outlined by Jo *et al.* (79, 81). Then, each system was simulated for 50 ns.

#### Derivation of depth-dependent water density profiles

MDAnalysis (82) was used to extract water density information from each lipid bilayer simulation trajectory. For each frame, the system was first re-centered on the lipid center-of-mass. Then, we computed a normalized histogram of TIP3 *z*-coordinates with 1Å bins to capture the distribution of water molecules. The time-averaged histogram was computed by averaging the histograms representing each frame (Fig. S2).

To generate analytic profiles, we used nonlinear regression to fit each histogram to the logistic function, *f*_thk_:

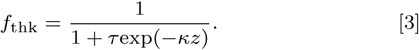

The function *f*_thk_ depends on membrane depth (*z*) and has two adjustable parameters: steepness *κ* and width *τ*. We derived *κ* and *τ* for all simulated lipid compositions. The resulting parameters are listed in Table S5 and the analytic water density profiles are shown in Fig. S3.

#### Calculation of water-filled pore shapes

For proteins with more than three transmembrane segments, we introduced a pore into the implicit membrane model. To determine the pore shape, we created a new method to transform discrete structural information into a smooth geometry described by differentiable functional forms. First, we used the convex-hull algorithm described in Koehler Leman *et al* (83) to identify backbone and side chain atoms that are in the transmembrane region (|*z*| *≤ T*), face the protein interior, and are not buried. A side-chain was defined as buried if it had 23 or more neighboring atoms within 12 Å of its *C*_*α*_ atom (59). Next, we computed a histogram of the *z*-coordinates of pore facing atoms with a bin size of 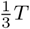. For each bin, the (*x, y*) coordinates of the atoms were collected. Then, the Khachiyan algorithm (84) was used to compute the minimum-area ellipse that bounds these coordinates. Each ellipse is defined with the following parameters: major radius (*a*), minor radius (*b*), rotation angle (*θ*), and center (*x*_0_,*y*_0_). The radius of the ellipse, *g*_radius_, is calculated using rotation matrix *M*:

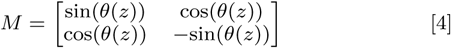

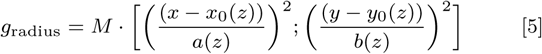

Cubic spline interpolation was used to fit polynomials to describe the depth-dependence of each parameter. The result is five continuous and differentiable parametric functions: *a*(*z*), *b*(*z*),*θ*(*z*), *x*_0_(*z*), and *y*_0_(*z*). The transition between the water-filled pore and lipid phase is defined by *g*_radius_ given the transition steepness *n*:

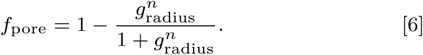

#### Validation of model parameters

##### 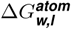 values

To verify 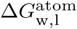 values, we first recalculated the side chain transfer energies by solving **Ax** = **b**. The Pearson correlation coefficient between the calculated and experimentally measured side chain transfer energies was *R*^2^ = 0.99 (Fig. S1). In addition, we used the procedure outlined in the *Scientific Benchmarks* section of *Methods* to estimate the ΔΔ*G*_mut_ values from Moon & Fleming (45). Specifically, we sought to verify that 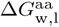 trends were preserved in context of the full energy function. The correlation between predicted and experimentally measured ΔΔ*G*^mut^ values was R^2^ = 0.84. Note, the ΔΔ*G*^mut^ for proline was excluded from the correlation coefficients because steric clashes resulted large energies.

##### Membrane thickness

We validated the water-density profiles computed from Molecular Dynamics simulations by comparing the derived membrane thickness parameters with thickness measured at various temperatures via x-ray and neutron scattering experiments (85–87). First, we computed the membrane half thickness *t* from each logistic curve as the Gibbs dividing surface between the water and lipid phases (*f* (*z*) = 0.5). We then calculated a line of best fit through the experimental measured thickness values at each temperature (Fig. S4).

#### Evaluation through scientific benchmarks

##### Test #1: Prediction of transmembrane helical peptide orientation

Low-free-energy peptide orientations were identified by calculating an energy landscape: a mapping between all possible peptide orientations relative to the membrane and their energies. Orientation was defined by two coordinates: (1) distance between the membrane center and peptide center of mass, *d* and (2) angle between the membrane normal and helical axis, *θ*. To compute the mapping, we first applied side-chain packing and minimization to resolve steric clashes in the peptide structure. Then, we applied rigid-body moves to sample all combinations of *θ* and *d* values. Membrane depths were sampled between −60 Å and 60 Å with a 1 Å step size and tilt angles were sampled between 0° −360° with a 1°step size. In addition, the water-to-bilayer transfer energy Δ*G*_w,l_ was computed for each peptide as the difference in energy between the aqueous phase (Δ*G*_w_: peptide at (60 Å,270 °)) and the lipid phase (Δ*G*_l_: peptide at (0 Å, 270°)): Δ*G*_w,l_ = Δ*G*_l_ - Δ*G*_w_.

We computed energy landscapes for five transmembrane helical peptides (53): acetylcholine receptor segment (1a11), M2 proton channel segment (1mp6) NMDA glutamate receptor (2nr1), VPU-forming domain of HIV-1 (1pje), and WALP23. Coordinates for the first four peptides were downloaded from the Protein Data Bank (88). The structure of WALP23 was modeled as an ideal helix with *ϕ* = −47°and *ψ* = −57°. For calculations with the new implicit membrane model, we chose lipid composition parameters that were consistent with the lipid composition used for the experimental measurement (Table S8).

##### Test #2: ΔΔ*G*^*mut*^predictions

ΔΔ*G*^mut^ values were computed using the protocol described in Alford *et al.* (57). Here, a mutation is introduced at the host site and the side chains are optimized within 8Å of the mutated residue. The ΔΔ*G*^mut^ was calculated as the difference in energy between the mutant (Δ*G*^mutant^) and native (Δ*G*^native^) conformations: ΔΔ*G*^mut^ = Δ*G*^mutant^ - Δ*G*^native^. ΔΔ*G*^mut^ prediction was evaluated on mutations in position 111 in outer membrane palmitoyl transferase (PagP; 3qd6) (56). The dataset included mutations from the native amino acid to all 19 other canonical amino acids. For calculations with the new implicit membrane model, parameters for DLPC membranes at 20° were chosen to match experimental conditions.

##### Test # 3: Native structure discrimination

Native structure discrimination is the ability of an energy function to distinguish near-native from non-native conformations. To measure discrimination, we used ensembles of five *α*-helical proteins generated by Dutagaci *et al.* (58): bacteriorhodopsin (BRD7; 1py6), fumarate reductase (FMR5; 1qla), lactose permease (LTPA; 1pv6), rhodopsin (RHOD; 1u19), and V-ATPase (VATP; 2bl2). Structures in each ensemble were between 1-11Å from the coordinates of the crystal structure. We refined each conformation using RosettaMPRelax (57) with constraints to the starting *C*_*α*_ coordinates. Then, we computed the Boltzmann-weighted average root-mean-squared-deviation (RMS) *W*_RMS_ (89). Here, *W*_RMS_ is computed over *N* refined candidates with scores Δ*G* and distances from the native RMS at a given temperature *T* (Eq. 7):

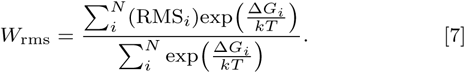

##### Test #4: Native sequence recovery

The fourth benchmark test was sequence recovery: the fraction of amino acids recovered after performing complete redesign of naturally occurring proteins. To evaluate recovery, we used a benchmark set of 133 *α*-helical and *β*-barrel proteins, each between 40-10,000 residues with less than 25% sequence identity and better than 3.0 Å resolution (83). The protein coordinates were downloaded from the Orientations of Proteins in Membranes database (90). Then, the fixed-backbone Rosetta Design protocol (91) was used to search for low-free energy-sequences. Sequence recovery, *R*_seq_, was calculated as the number of correct positions *N*_correct_ relative to the number of available positions *N*_all_:

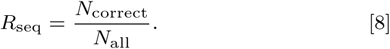

In addition, we examined sequence recovery rates for individual amino acid types, relative the background probability of guessing a random amino acid type (1 in 20 types, or 5%). This metric, *N*_AA_ is the fraction of amino acid types recovered with rates higher than random.

##### Properties of *in silico* designs

We examined properties of the *in silico* designed proteins using the Kullback-Leibler (KL) divergence. This metric quantifies the divergence between the distribution of amino acid types (*i*) in the native (*N*_nat,*i*_) and designed (*N*_des,*i*_) sequences:

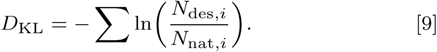

##### Data Availability

The energy function and benchmark methods available in the the Rosetta Software Suite. Rosetta is available to non-commercial users for free and to commercial users for a fee. The detailed command lines are provided in the Supplementary Information.

## ACKNOWLEDGMENTS

Thank you to Sergey Lyskov, Vikram Mulligan, and Julia Koehler for code reviews. We also thank Sai Pooja Mahajan for critical reading of the manuscript. R.F.A. was supported by a Hertz Foundation Fellowship and a National Science Foundation Graduate Research Fellowship. This work was also supported by National Institute of Health Grants GM-072281 (R.F.A. and J.J.G.) and GM-079440 (P.J.F. and K.G.F.). Computations were performed using the Maryland Advanced Research Computing Center and the National Science Foundation XSEDE Grant TG-MCB180056.

Conceptualization & Methodology, R.F.A, P.J.F., K.G.F., and J.J.G., Investigation, Data Curation, Software, and Analysis, R.F.A., Writing - Original Draft, R.F.A., Writing - Review & Editing, R.F.A., P.J.F., K.G.F., and J.J.G., Funding Acquisition, R.F.A., K.G.F., and J.J.G., Resources & Supervision, K.G.F., J.J.G.

## The authors declare the following competing financial interests

Dr. Gray is an unpaid board member of the Rosetta Commons. Under institutional participation agreements between the University of Washington, acting on behalf of the Rosetta Commons, Johns Hopkins University may be entitled to a portion of revenue received on licensing Rosetta software including programs described here. As a member of the Scientific Advisory Board of Cyrus Biotechnology, JJG is granted stock options. Cyrus Biotechnology distributes the Rosetta software, which may include methods described in this paper.

## References

1. Huang PS, Boyken SE, Baker D (2016) The coming of age of de novo protein design. Nature 537(7620):320–327.

2. Samish I, MacDermaid CM, Perez-Aguilar JM, Saven JG (2011) Theoretical and Computational Protein Design. Annual Review of Physical Chemistry 62(1):129–149.

3. Barth P, Senes A (2016) Toward high-resolution computational design of the structure and function of helical membrane proteins. Nature Structural & Molecular Biology 23(6):475–480.

4. Tan S, Tan HT, Chung MCM (2008) Membrane proteins and membrane proteomics. PROTEOMICS 8(19):3924–3932.

5. Overington JP, Al-Lazikani B, Hopkins AL (2006) How many drug targets are there? Nature Reviews Drug Discovery 5(12):993–996.

6. Joh NH, et al. (2014) De novo design of a transmembrane Zn 2+-transporting four-helix bundle. Science 346(6216):1520–1524.

7. Mahendran KR, et al. (2017) A monodisperse transmembrane *α*-helical peptide barrel. Nature Chemistry 9(5):411–419.

8. Chowdhury R, et al. (2018) PoreDesigner for tuning solute selectivity in a robust and highly permeable outer membrane pore. Nature Communications 9(1):3661.

9. Young M, et al. (2018) Computational design of orthogonal membrane receptor-effector switches for rewiring signaling pathways. Proceedings of the National Academy of Sciences 115(27):7051–7056.

10. Feng X, Ambia J, Chen KYM, Young M, Barth P (2017) Computational design of ligand-binding membrane receptors with high selectivity. Nature Chemical Biology 13(7):715–723.

11. Lu P, et al. (2018) Accurate computational design of multipass transmembrane proteins. Science (New York, N.Y.) 359(6379):1042–1046.

12. Dickson CJ, Rosso L, Betz RM, Walker RC, Gould IR (2012) GAFFlipid: a General Amber Force Field for the accurate molecular dynamics simulation of phospholipid. Soft Matter 8(37):9617.

13. Klauda JB, et al. (2010) Update of the CHARMM All-Atom Additive Force Field for Lipids: Validation on Six Lipid Types. The Journal of Physical Chemistry B 114(23):7830–7843.

14. Venable RM, Brown FL, Pastor RW (2015) Mechanical properties of lipid bilayers from molecular dynamics simulation. Chemistry and Physics of Lipids 192:60–74.

15. Jämbeck JPM, Lyubartsev AP (2012) Derivation and Systematic Validation of a Refined All-Atom Force Field for Phosphatidylcholine Lipids. The Journal of Physical Chemistry B 116(10):3164–3179.

16. Schmid N, et al. (2011) Definition and testing of the GROMOS force-field versions 54A7 and 54B7. European Biophysics Journal 40(7):843–856.

17. Leonard AN, Wang E, Monje-Galvan V, Klauda JB (2019) Developing and Testing of Lipid Force Fields with Applications to Modeling Cellular Membranes. Chemical Reviews p. acs.chemrev.8b00384.

18. Phillips JC, Sun Y, Jain N, Bohm EJ, Kale LV (2014) Mapping to Irregular Torus Topologies and Other Techniques for Petascale Biomolecular Simulation in SC14: International Conference for High Performance Computing, Networking, Storage and Analysis. (IEEE), pp. 81–91.

19. Marrink SJ, et al. (2019) Computational Modeling of Realistic Cell Membranes. Chemical Reviews p. acs.chemrev.8b00460.

20. Marrink SJ, Risselada HJ, Yefimov S, Tieleman DP, de Vries AH (2007) The MARTINI Force Field: Coarse Grained Model for Biomolecular Simulations. The Journal of Physical Chemistry B 111(27):7812–7824.

21. Orsi M, Essex JW (2011) The ELBA Force Field for Coarse-Grain Modeling of Lipid Membranes. PLoS ONE 6(12):e28637.

22. Barrera EE, Frigini EN, Porasso RD, Pantano S (2017) Modeling DMPC lipid membranes with SIRAH force-field. Journal of Molecular Modeling 23(9):259.

23. Baoukina S, Rozmanov D, Tieleman DP (2017) Composition Fluctuations in Lipid Bilayers. Biophysical Journal 113(12):2750–2761.

24. Warshel A, Russell ST (1984) Calculations of electrostatic interactions in biological systems and in solutions. Quarterly Reviews of Biophysics 17(03):283.

25. Grossfield A (2008) Chapter 5 Implicit Modeling of Membranes. Current Topics in Membranes 60:131–157.

26. Ulmschneider JP, Ulmschneider MB (2009) Sampling efficiency in explicit and implicit membrane environments studied by peptide folding simulations. Proteins: Structure, Function, and Bioinformatics 75(3):586–597.

27. Davis ME, McCammon JA (1990) Electrostatics in biomolecular structure and dynamics. Chemical Reviews 90(3):509–521.

28. Baker NA, Sept D, Joseph S, Holst MJ, McCammon JA (2001) Electrostatics of nanosystems: application to microtubules and the ribosome. Proceedings of the National Academy of Sciences of the United States of America 98(18):10037–41.

29. Choe S, Hecht KA, Grabe M (2008) A continuum method for determining membrane protein insertion energies and the problem of charged residues. The Journal of general physiology 131(6):563–73.

30. Im W, Lee MS, Brooks CL (2003) Generalized born model with a simple smoothing function. Journal of Computational Chemistry 24(14):1691–1702.

31. Spassov V, Yan L, Sándor S (2002) Introducing an Implicit Membrane in Generalized Born/Solvent Accessibility Continuum Solvent Models. The Journal of Physical Chemistry B 106(34):8726–8738.

32. Tsui V, Case DA (2000) Theory and applications of the generalized born solvation model in macromolecular simulations. Biopolymers 56(4):275–291.

33. Panahi A, Feig M (2013) Dynamic Heterogeneous Dielectric Generalized Born (DHDGB): An Implicit Membrane Model with a Dynamically Varying Bilayer Thickness. Journal of Chemical Theory and Computation 9(3):1709–1719.

34. Lazaridis T, Karplus M (1999) Effective energy function for proteins in solution. Proteins 35(2):133–52.

35. Lazaridis T (2003) Effective energy function for proteins in lipid membranes. Proteins: Structure, Function, and Genetics 52(2):176–192.

36. Radzicka A, Wolfenden R (1988) Comparing the polarities of the amino acids: side-chain distribution coefficients between the vapor phase, cyclohexane, 1-octanol, and neutral aqueous solution. Biochemistry 27(5):1664–1670.

37. Nepal B, Leveritt J, Lazaridis T (2018) Membrane Curvature Sensing by Amphipathic Helices: Insights from Implicit Membrane Modeling. Biophysical Journal 114(9):2128–2141.

38. Yarov-Yarovoy V, Schonbrun J, Baker D (2005) Multipass membrane protein structure prediction using Rosetta. Proteins: Structure, Function, and Bioinformatics 62(4):1010–1025.

39. Barth P, Schonbrun J, Baker D (2007) Toward high-resolution prediction and design of transmembrane helical protein structures. Proceedings of the National Academy of Sciences 104(40):15682–15687.

40. Franks NP, Abraham MH, Lieb WR (1993) Molecular organization of liquid n-octanol: an X-ray diffraction analysis. Journal of pharmaceutical sciences 82(5):466–70.

41. White S (2018) Membrane Proteins of Known Structure.

42. MacCallum JL, Bennett WFD, Tieleman DP (2008) Distribution of amino acids in a lipid bilayer from computer simulations. Biophysical journal 94(9):3393–404.

43. Koehler Leman J, Bonneau R, Ulmschneider MB (2018) Statistically derived asymmetric membrane potentials from *α*-helical and *β*-barrel membrane proteins. Scientific Reports 8(1):4446.

44. Senes A, et al. (2007) Ez, a Depth-dependent Potential for Assessing the Energies of Insertion of Amino Acid Side-chains into Membranes: Derivation and Applications to Determining the Orientation of Transmembrane and Interfacial Helices. Journal of Molecular Biology 366(2):436–448.

45. Moon CP, Fleming KG (2011) Side-chain hydrophobicity scale derived from transmembrane protein folding into lipid bilayers. Proceedings of the National Academy of Sciences 108(25):10174–10177.

46. Harayama T, Riezman H (2018) Understanding the diversity of membrane lipid composition. Nature Reviews Molecular Cell Biology 19(5):281–296.

47. Lazaridis T (2005) Structural Determinants of Transmembrane *β*-Barrels. Journal of Chemical Theory and Computation 1(4):716–722.

48. Pellegrini-Calace M, Maiwald T, Thornton JM (2009) PoreWalker: A Novel Tool for the Identification and Characterization of Channels in Transmembrane Proteins from Their Three-Dimensional Structure. PLoS Computational Biology 5(7):e1000440.

49. Alford RF, et al. (2017) The Rosetta All-Atom Energy Function for Macromolecular Modeling and Design. Journal of Chemical Theory and Computation 13(6):3031–3048.

50. Yarov-Yarovoy V, et al. (2012) Structural basis for gating charge movement in the voltage sensor of a sodium channel. Proceedings of the National Academy of Sciences of the United States of America 109(2):E93–102.

51. Park H, et al. (2016) Simultaneous Optimization of Biomolecular Energy Functions on Features from Small Molecules and Macromolecules. Journal of Chemical Theory and Computation 12(12):6201–6212.

52. Killian J (2003) Synthetic peptides as models for intrinsic membrane proteins. FEBS Letters 555(1):134–138.

53. Ulmschneider MB, Sansom MSP, Di Nola A (2006) Evaluating tilt angles of membrane-associated helices: comparison of computational and NMR techniques. Biophysical journal 90(5):1650–60.

54. Kim T, Im W (2010) Revisiting Hydrophobic Mismatch with Free Energy Simulation Studies of Transmembrane Helix Tilt and Rotation. Biophysical Journal 99(1):175–183.

55. Strandberg E, et al. (2004) Tilt angles of transmembrane model peptides in oriented and non-oriented lipid bilayers as determined by 2H solid-state NMR. Biophysical journal 86(6):3709–21.

56. Marx DC, Fleming KG (2017) Influence of Protein Scaffold on Side-Chain Transfer Free Energies. Biophysical journal 113(3):597–604.

57. Alford RF, et al. (2015) An Integrated Framework Advancing Membrane Protein Modeling and Design. PLOS Computational Biology 11(9):e1004398.

58. Dutagaci B, Wittayanarakul K, Mori T, Feig M (2017) Discrimination of Native-like States of Membrane Proteins with Implicit Membrane-based Scoring Functions. Journal of Chemical Theory and Computation 13(6):3049–3059.

59. Kuhlman B, Baker D (2000) Native protein sequences are close to optimal for their structures. Proceedings of the National Academy of Sciences of the United States of America 97(19):10383–8.

60. Wallin E, Tsukihara T, Yoshikawa S, Heijne GV, Elofsson A (2008) Architecture of helix bundle membrane proteins: An analysis of cytochrome c oxidase from bovine mitochondria. Protein Science 6(4):808–815.

61. McDonald SK, Fleming KG (2016) Aromatic Side Chain Water-to-Lipid Transfer Free Energies Show a Depth Dependence across the Membrane Normal. Journal of the American Chemical Society 138(25):7946–7950.

62. Lazaridis T (2004) Implicit solvent simulations of peptide interactions with anionic lipid membranes. Proteins: Structure, Function, and Bioinformatics 58(3):518–527.

63. Lazaridis T, Leveritt JM, PeBenito L (2014) Implicit membrane treatment of buried charged groups: Application to peptide translocation across lipid bilayers. Biochimica et Biophysica Acta (BBA) -Biomembranes 1838(9):2149–2159.

64. Rothman JE, Lenard J (1977) Membrane asymmetry. Science (New York, N.Y.) 195(4280):743–53.

65. Jarsch IK, Daste F, Gallop JL (2016) Membrane curvature in cell biology: An integration of molecular mechanisms. The Journal of cell biology 214(4):375–87.

66. Lai JK, Ambia J, Wang Y, Barth P (2017) Enhancing Structure Prediction and Design of Soluble and Membrane Proteins with Explicit Solvent-Protein Interactions. Structure 25(11):1758–1770.e8.

67. Vermaas JV, Pogorelov TV, Tajkhorshid E (2017) Extension of the Highly Mobile Membrane Mimetic to Transmembrane Systems through Customized in Silico Solvents. The journal of physical chemistry. B 121(15):3764–3776.

68. MacKerell AD, et al. (1998) All-Atom Empirical Potential for Molecular Modeling and Dynamics Studies of Proteins <sup>†</sup>. The Journal of Physical Chemistry B 102(18):3586–3616.

69. Ponder JW, Case DA (2003) Force fields for protein simulations. Advances in protein chemistry 66:27–85.

70. Jorgensen WL, Maxwell DS, Tirado-Rives J (1996) Development and Testing of the OPLS All-Atom Force Field on Conformational Energetics and Properties of Organic Liquids. Journal of the American Chemical Society.

71. O’Meara MJ, et al. (2015) Combined Covalent-Electrostatic Model of Hydrogen Bonding Improves Structure Prediction with Rosetta. Journal of Chemical Theory and Computation 11(2):609–622.

72. Yuzlenko O, Lazaridis T (2013) Membrane protein native state discrimination by implicit membrane models. Journal of computational chemistry 34(9):731–8.

73. Dutagaci B, Feig M (2017) Determination of Hydrophobic Lengths of Membrane Proteins with the HDGB Implicit Membrane Model. Journal of Chemical Information and Modeling 57(12):3032–3042.

74. Kroncke BM, et al. (2016) Documentation of an Imperative To Improve Methods for Predicting Membrane Protein Stability. Biochemistry 55(36):5002–9.

75. Duran AM, Meiler J (2018) Computational design of membrane proteins using RosettaMembrane. Protein Science 27(1):341–355.

76. Mobley DL, et al. (2018) Escaping Atom Types in Force Fields Using Direct Chemical Perception. Journal of Chemical Theory and Computation 14(11):6076–6092.

77. Lamy-Freund MT, Riske KA (2003) The peculiar thermo-structural behavior of the anionic lipid DMPG. Chemistry and physics of lipids 122(1-2):19–32.

78. Leonenko ZV, Finot E, Ma H, Dahms TES, Cramb DT (2004) Investigation of temperature-induced phase transitions in DOPC and DPPC phospholipid bilayers using temperature-controlled scanning force microscopy. Biophysical journal 86(6):3783–93.

79. Wu EL, et al. (2014) CHARMM-GUI Membrane Builder toward realistic biological membrane simulations. Journal of Computational Chemistry 35(27):1997–2004.

80. Phillips JC, et al. (2005) Scalable molecular dynamics with NAMD. Journal of Computational Chemistry 26(16):1781–1802.

81. Jo S, Lim JB, Klauda JB, Im W (2009) CHARMM-GUI Membrane Builder for Mixed Bilayers and Its Application to Yeast Membranes. Biophysical Journal 97(1):50–58.

82. Michaud-Agrawal N, Denning EJ, Woolf TB, Beckstein O (2011) MDAnalysis: a toolkit for the analysis of molecular dynamics simulations. Journal of computational chemistry 32(10):2319–27.

83. Koehler Leman J, Lyskov S, Bonneau R (2017) Computing structure-based lipid accessibility of membrane proteins with mp_lipid_acc in RosettaMP. BMC Bioinformatics 18(1):115.

84. Khachiyan LG, Todd MJ (1993) On the complexity of approximating the maximal inscribed ellipsoid for a polytope. Mathematical Programming 61(1-3):137–159.

85. Kučerka N, Nieh MP, Katsaras J (2011) Fluid phase lipid areas and bilayer thicknesses of commonly used phosphatidylcholines as a function of temperature. Biochimica et Biophysica Acta (BBA) - Biomembranes 1808(11):2761–2771.

86. Kučerka N, Holland BW, Gray CG, Tomberli B, Katsaras J (2012) Scattering Density Profile Model of POPG Bilayers As Determined by Molecular Dynamics Simulations and Small-Angle Neutron and X-ray Scattering Experiments. The Journal of Physical Chemistry B 116(1):232–239.

87. Kučerka N, et al. (2015) Molecular Structures of Fluid Phosphatidylethanolamine Bilayers Obtained from Simulation-to-Experiment Comparisons and Experimental Scattering Density Profiles. The Journal of Physical Chemistry B 119(5):1947–1956.

88. Berman HM, et al. (2000) The Protein Data Bank. Nucleic acids research 28(1):235–42.

89. Bhardwaj G, et al. (2016) Accurate de novo design of hyperstable constrained peptides. Nature 538(7625):329–335.

90. Lomize MA, Pogozheva ID, Joo H, Mosberg HI, Lomize AL (2012) OPM database and PPM web server: resources for positioning of proteins in membranes. Nucleic Acids Research 40(D1):D370–D376.

91. Leaver-Fay A, et al. (2013) Scientific benchmarks for guiding macromolecular energy function improvement. Methods in enzymology 523:109–43.

