## Supporting Information for "Protein structure prediction and design in a biologically-realistic implicit membrane"

#### This PDF file includes:

Supplementary text
Figs. S1 to S7
Tables S1 to S14

### Supporting Information Text

### Detailed methods and command lines

**A. Benchmark #1: Membrane energy landscapes.** Benchmark #1 traverses the macromolecular energy landscape of transmembrane peptides as a function of protein orientation. The peptide is first transformed into the membrane coordinate frame. This step is followed by side-chain repacking and energy minimization of all torsion angles ($\phi$, $\psi$, and $\omega$). Then, the `MembraneEnergyLandscapeSampler` samples and scores all peptide orientations between $-60$Å and 60Å with a 1Å step size and $0 - 360°$ with a 1° step size. The protocol is captured by the XML script listed below.

```
<ROSETTASCRIPTS>
        <SCOREFXNS>
                <ScoreFunction name="s" weights="%%sfxn_weights%%"/>
        </SCOREFXNS>
        <TASKOPERATIONS>
                <RestrictToRepacking name="rtrp"/>
                <ExtraRotamersGeneric name="extra_chi" ex1="1"
                ex2="1" extrachi_cutoff="0"/>
        </TASKOPERATIONS>
        <MOVERS>
                <AddMembraneMover name="add_memb" />
                <TransformIntoMembraneMover name="transform_into_memb" />
                <PackRotamersMover name="pack_rotamers" scorefxn="s"
                task_operations="rtrp"/>
                <MinMover name="minimize_struc" scorefxn="s" chi="1" bb="1" jump="0"
                type="dfpmin_armijo_nonmonotone" tolerance="0.01" />
                <MembraneEnergyLandscapeSampler name="landscape_test"
                scorefxn="s" interface="0" />
        </MOVERS>
        <PROTOCOLS>
                <Add mover_name="add_memb"/>
                <Add mover_name="transform_into_memb"/>
                <Add mover_name="pack_rotamers"/>
                <Add mover_name="minimize_struc"/>
                <Add mover_name="landscape_test"/>
        </PROTOCOLS>
</ROSETTASCRIPTS>
```

The protocol is deterministic and outputs a single file with a mapping between orientations and energies. To run the protocol, we ran the following command line for each peptide:

```
/path/to/Rosetta/main/source/bin/rosetta_scripts.macosclangrelease
    -in:file:s 1a11.pdb                            # Input PDB File
    -mp:setup:spanfiles 1a11.span                  # Input span file
    -parser:protocol sample_energy_landscape.xml   # Name of XML script file
    -parser:script_vars
        sfxn_weights=franklin2019                  # Energy function to use
    -mp:lipids:composition DLPC                    # Lipid composition (for franklin2019)
```

**B. Benchmark #2: $\Delta\Delta G^{\mathrm{mut}}$ prediction.** To calculate $\Delta\Delta G_{\mathrm{mut}}$ values, we used the MPddG protocol described in Alford *et al.* (? ). The protocol combines the RosettaMP framework with a fixed-backbone $\Delta\Delta G$ prediction protocol similar to the method described in Kellogg *et al.* (? ). For each $\Delta\Delta G$ prediction, a new side chain was introduced at the host site and side-chains within 8Å were repacked. Then, the $\Delta\Delta G$ was calculated as the difference between the mutant and native conformation energies in Rosetta Energy Units (REU).

The MPddG protocol is captured in a PyRosetta script (? ). The following command line and options were used for prediction of $\Delta\Delta G_{\mathrm{mut}}$ values for mutations in the OmpLA and PagP scaffolds:

```
python predict_ddG.py
   --pdb sample.pdb               # Input PDB file
   --spanfile sample.span         # Input spanfile
   --out_ddGs ddGs.txt            # Name of file with predicted ddGs
   --out_breakdown decomposed.sc  # Name of file with predicted per-term ddGs
   --lipid_composition DLPC       # Lipid composition parameters to use
   --res ##                       # Host site (181 for OmpLA, 104 for PagP)
   --repack_radius 8.0            # Repack all residues within 8.0 \AA of host site
```

The script will predict $\Delta\Delta G_{\mathrm{mut}}$ values for all canonical amino acids. The first output file, `ddGs.txt` includes the predicted $\Delta\Delta G_{\mathrm{mut}}$ values. The second output file, `decomposed.sc` includes the contribution of each energy term to the overall $\Delta\Delta G_{\mathrm{mut}}$.

**C. Benchmark #3: Native structure discrimination against decoys.** To measure native structure discrimination, we refined decoys using the `mp_relax` protocol (? ). The `mp_relax` protocol combines the RosettaMP Framework with FastRelax (? ): a protocol that perturbs the protein using small backbone torsion moves, followed by side-chain repacking and energy minimization along all torsion angles ($\phi$, $\psi$, $\omega$). The protocol is captured by the Rosetta XML script `mp_relax.xml` listed below:

```
<ROSETTASCRIPTS>
        <SCOREFXNS>
                <ScoreFunction name="memb_hires" weights="%%sfxn_weights%%" />
        </SCOREFXNS>
        <MOVERS>
                <AddMembraneMover name="add_memb"/>
                <MembranePositionFromTopologyMover name="init_pos"/>
                <FastRelax name="fast_relax" scorefxn="memb_hires" repeats="8"/>
        </MOVERS>
        <PROTOCOLS>
                <Add mover="add_memb"/>
                <Add mover="init_pos"/>
                <Add mover="fast_relax"/>
        </PROTOCOLS>
        <OUTPUT scorefxn="memb_hires" />
</ROSETTASCRIPTS>
```

To execute the program, we ran the following command line on the decoy set for each target:

```
/path/to/Rosetta/main/source/bin/rosetta_scripts.linuxgccrelease
    -in:file:native sample_native.pdb   # Native PDB coordinates
    -in:file:l sample_candidates.list   # List of candidate models
    -mp:setup:spanfiles sample.span     # Path to spanning topology file
    -parser:script_vars
        sfxn_weights=franklin2019       # Name of candidate energy function
    -parser:protocol mp_relax.xml       # Path to XML script for relax protocol
    -out:file:scorefile model_scores.sc # Path to file with energies and RMS values
    -out:path:all /path/to/output/pdbs  # Path to refined models
    -mp:lipids:composition DLPC         # Choose phospholipid composition
```

To compute the RMSD between the native and refined models, we used the `score_jd2` application with the following options:

```
/path/to/Rosetta/main/source/bin/score_jd2.linuxgccrelease
    -in:file:l refined_models.list    # List of refined models
    -in:file:native native.pdb        # Native structure for RMSD calculation
    -in:file:spanfile sample.span     # Input spanfile
    -in:membrane                      # Use the RosettaMP Framework
    -score:weights franklin2019       # Use hi-res membrane score function
    -mp:lipid:composition DLPC        # Chose phospholipid composition
```

Finally, we estimated the native structure discrimination score, $W_{\mathrm{rms}}$ using `score_energy_landscape.py`:

```
/path/to/Rosetta/bakerlab_scripts/boinc/score_energy_landscape.py
    -terms rms total_score              # Name of rms and energy term
    -abinitio_scorefile candidates.sc   # Path to file with energies and rms values
```

**D. Benchmark #4: Protein Design.** To measure native sequence recovery, we used an adapted version of the fixed-backbone design protocol from Leaver-Fay *et al.* (REF). Here, we combined this protocol with the RosettaMP framework to keep proteins oriented in the bilayer during design. Each protein in our dataset was redesigned using the following command line:

```
/path/to/Rosetta/main/source/bin/fixbb.linuxgccrelease
    -in:file:s sample.pdb               # Input PDB coordinates
    -mp:setup:spanfiles sample.span     # Input spanfile
    -score:weights candidate_efxn       # Name of energy function weights file
```

```
    -in:membrane                        # Load the membrane framework
    -out:path:all                       # Path to output redesigned structures
    -in:ignore_unrecognized_res         # Ignore unknown ligand residues
    -mp:lipids:composition DOPC         # For M19 Only: Phospholipid composition
```

Then, we computed sequence recovery with the `mp_seqrecov` application using the command line given below:

```
/path/to/Rosetta/main/source/bin/mp_seqrecov.linuxgccrelease
    -native_pdb_list natives.list       # List of native protein PDBs
    -redesign_pdb_list redesigned.list  # List of redesigned protein PDBs
    -seq_recov_filename seqrecov.txt    # File for sequence recovery data
```

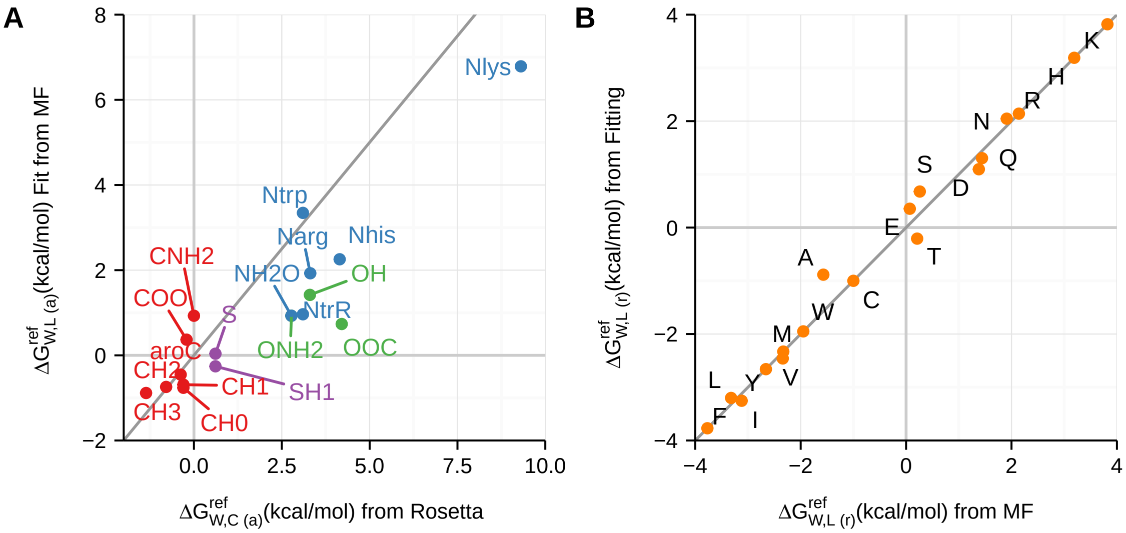


**Fig. S1.** Comparison of derived per-atom transfer energies for *franklin2019* with prior computational models (**M07**/**IMM1**) and experimental measurements. (A) Comparison of derived per-atom transfer energies with values from the Implicit Membrane Model 1. Carbon atom types are in red, nitrogen in blue, oxygen in green, and sulfur in purple. The mean absolute error (MAE) between IMM1 and derived values is 1.20 kcal/mol. (B) Comparison of calculated per-residue transfer energies with experimental values from Moon & Fleming. The MAE between calculated and experimental values is 0.15 kcal/mol.

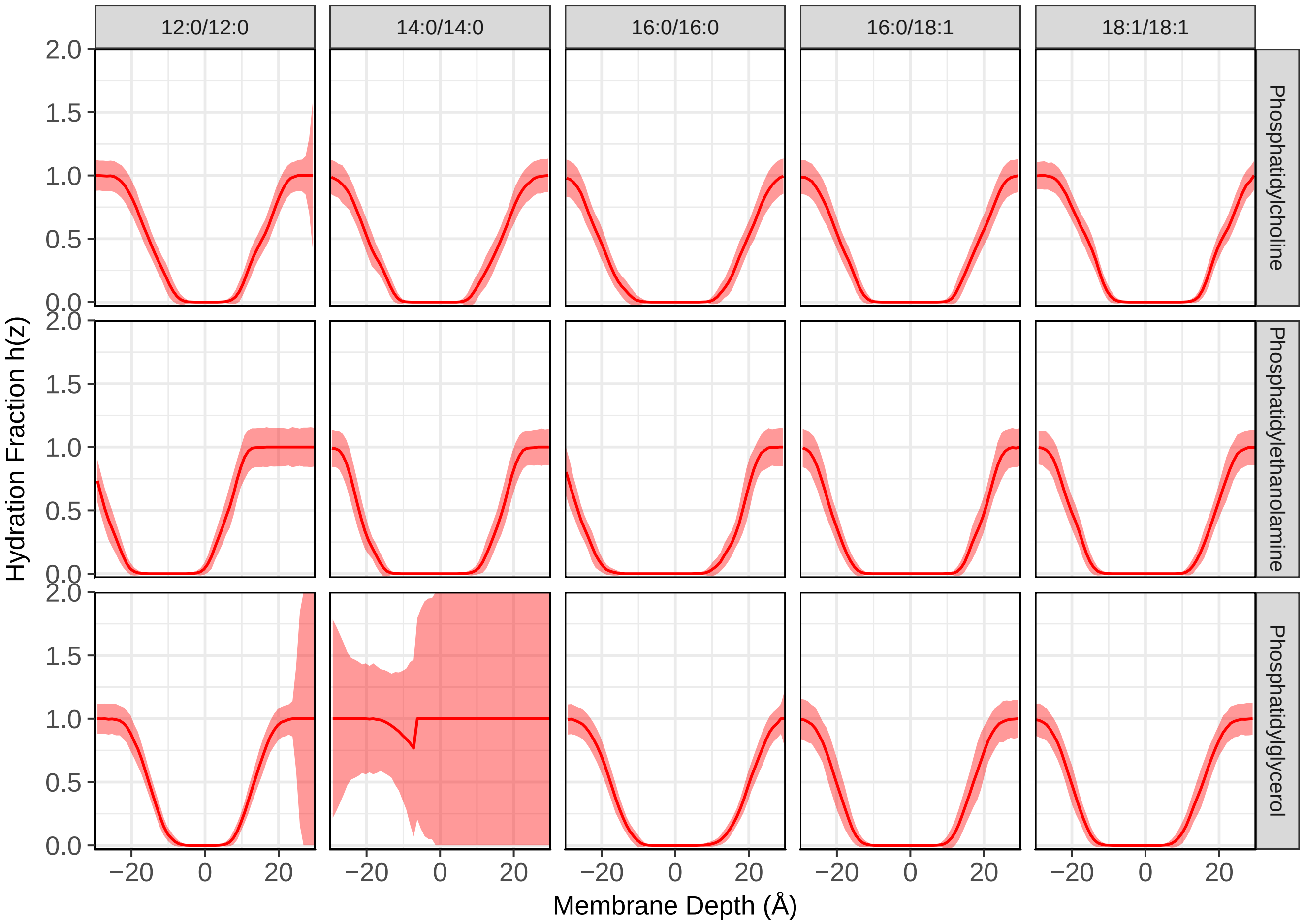


**Fig. S2.** Time-averaged density of water along the membrane normal axis for 15 lipid compositions. The time-averaged density is shown with a solid red line and the standard deviation across time steps is shaded in red.

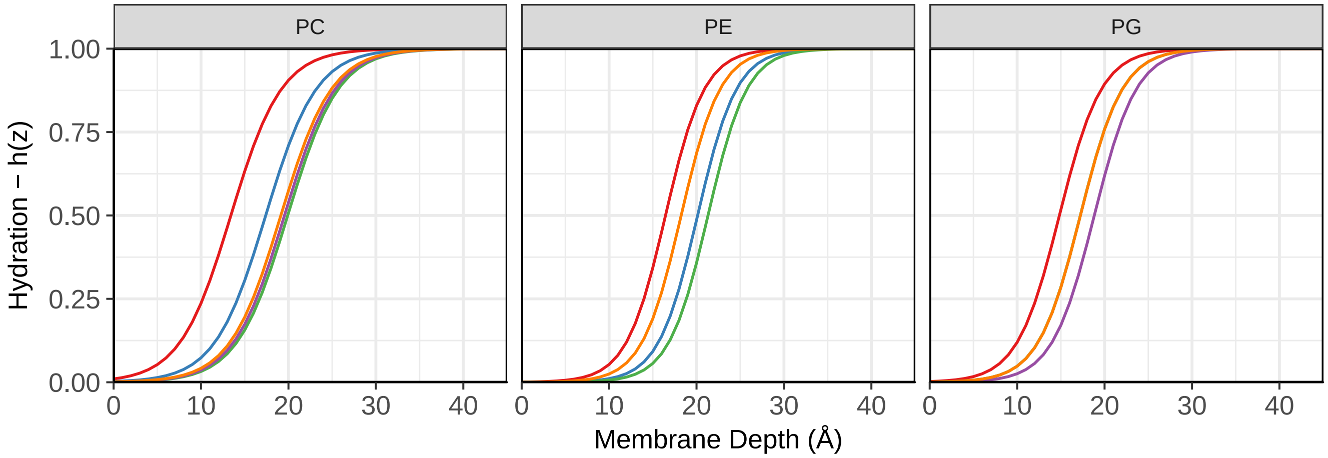


**Fig. S3.** Hydration profiles along the membrane normal for simulated phospholipid bilayers at physiological temperature. Each profile is colored by acyl chain type with 12:0/12:0 in red, 14:0/14:0 in blue, 16:0/16:0 in green, 18:1/18:1 in purple, and 16:0/18:1 in orange. For each lipid composition, membrane thickness increases with increasing acyl chain length. The exception is for 16:0/18:1 where the degree of saturation makes the chains more rigid and thins the bilayer.

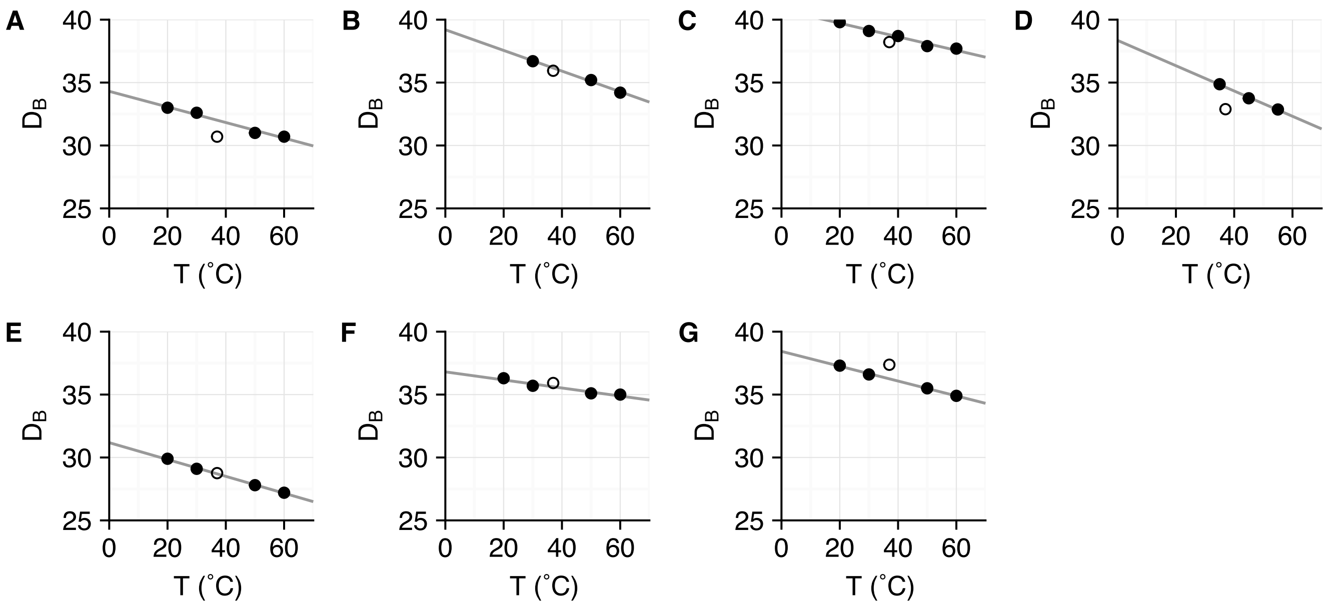


**Fig. S4.** Comparison of the membrane thickness computed from simulated water density and measured by X-Ray and neutron scattering experiments (? ? ? ? ). Measured thicknesses are represented by filled circles and thicknesses computed from molecular dynamics trajectories are represented by open circles. The solid line is the line of best fit through the measured values. (A) DLPC (B) DMPC (C) POPC (D) DLPE (E) DLPG (F) DOPG (G) POPG. The calculated values closely follow the thickness vs. temperature trend for each lipid composition.

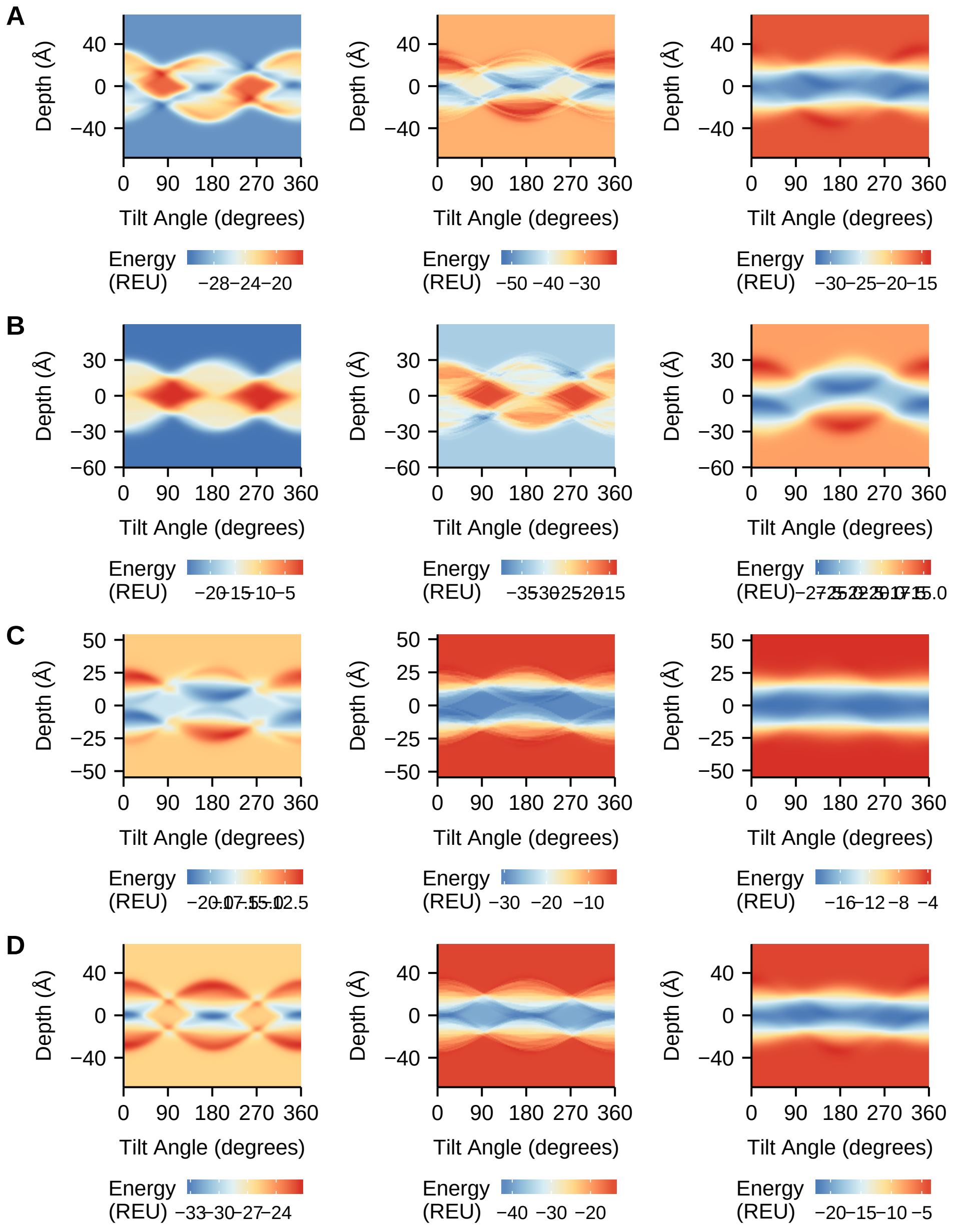


**Fig. S5.** Additional Protein energy landscapes for transmembrane $\alpha$-helical peptides. The energy landscape is calculated as a function of helix tilt angle and distance between the peptide center of mass and the membrane center. The points are colored by energy, with red representing a high energy conformation and blue representing a low energy conformation. (A) Energy landscape of $\alpha$-factor receptor M6 (1mp6) calculated by the **M07**, **M12**, and**M19M19** energy functions. (B) Energy landscape of VPU-forming domain of HIV-1 (1pje) calculated by the **M07**, **M12**, and**M19** energy functions. (C) Energy landscape of glutamate receptor NMDA subtype (2nr1) calculated by the **M07**, **M12**, and**M19** energy functions. (D) Energy landscape of the designed peptide WALP23 calculated by the **M07**, **M12**, and**M19** energy functions.

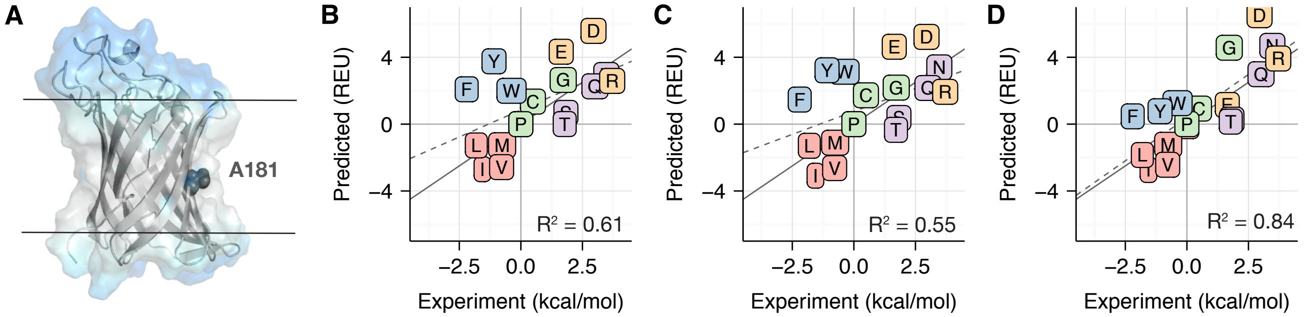


**Fig. S6.** Comparison between computationally predicted and experimentally measured $\Delta\Delta G_{\mathrm{mut}}$ values measured in the OmpLA scaffold (**?** ). For all correlation plots (B-D), proline is not shown due to steric clashes resulting in a large $\Delta\Delta G_{\mathrm{mut}}$ value. The dotted gray line is the line of best fit and the solid gray line is $y = x$. Amino acids are colored according to the following categories: charged (orange), nonpolar (red), aromatic (blue), polar (purple), special case (green). (A) Structure of the OmpLA scaffold (PDB 1qd6) with the mutation site A181 highlighted in dark grey. The implicit solvent phases are colored in a similar manner to Fig. 1 (Main Text). The $\Delta\Delta G_{\mathrm{mut}}$ predictions for mutations in OmpLA by M07, M12, and M19 are shown in panels B, C, and D respectively.

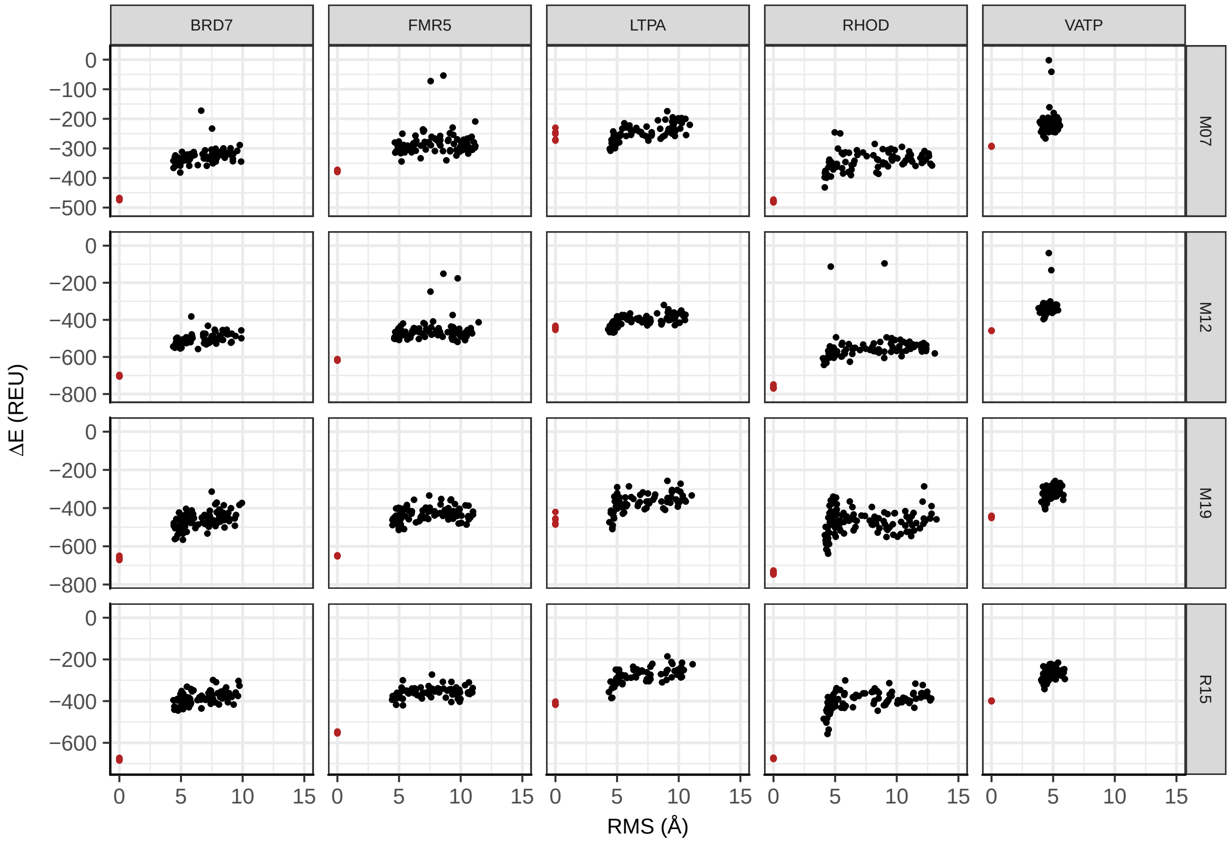


**Fig. S7.** Discrimination of target candidate structures from the native crystal structure. In each row, energy vs. RMS plots are shown for candidate models refined by the R15, M07, M12, and M19 energy functions. The refined candidate models are shown in black and refined natives are shown in red.

**Table S1. Rosetta atom types**

| Atom Type | Description | Atom Type | Description |
|---|---|---|---|
| CH0 | carbon with no hydrogens | CH2 | carbon with two hydrogens |
| CH1 | carbon with one hydrogen | CH3 | methyl carbon |
| CNH2 | Glu amine carbon | COO | carbonyl carbon |
| S | sulfur | SH1 | sulfur with one hydrogen |
| aroC | aromatic carbon | Nhis | His nitrogen |
| Ntrp | Trp and HIs nitrogen | Nlys | Lys nitrogen |
| NH2O | Glu amine nitrogen | Narg | arginine nitrogen |
| NtrR | Trp secondary nitrogen | ONH2 | Glu amine oxygen |
| OOC | carbonyl oxygen | OH | hydroxyl oxygen |

**Table S2. Constraint equations for water-to-bilayer transfer energy fit.**

| Amino Acid | Equation |
|---|---|
| A | $x_{\text{CH3}}$ |
| C | $x_{\text{CH2}} + x_{\text{SH1}}$ |
| D | $x_{\text{CH2}} + x_{\text{COO}} + 2x_{\text{OOC}}$ |
| E | $2x_{\text{CH2}} + x_{\text{COO}} + 2x_{\text{OOC}}$ |
| F | $x_{\text{CH0}} + x_{\text{CH2}} + 5x_{\text{aroC}}$ |
| G | – |
| H | $x_{\text{CH0}} + x_{\text{CH2}} + 2x_{\text{aroC}} + x_{\text{Nhis}} + x_{\text{Ntrp}}$ |
| I | $2x_{\text{CH3}} + 2x_{\text{CH2}}$ |
| K | $4x_{\text{CH2}} + x_{\text{Nlys}}$ |
| L | $2x_{\text{CH3}} + x_{\text{CH2}} + x_{\text{CH1}}$ |
| M | $x_{\text{CH3}} + 2x_{\text{CH2}} + x_{\text{S}}$ |
| N | $x_{\text{CH2}} + x_{\text{ONH2}} + x_{\text{CNH2}} + x_{\text{NH2O}}$ |
| P | $x_{\text{Npro}} + 3x_{\text{CH2}}$ |
| Q | $2x_{\text{CH2}} + x_{\text{ONH2}} + x_{\text{CNH2}} + x_{\text{NH2O}}$ |
| R | $3x_{\text{CH2}} + x_{\text{aroC}} + 2x_{\text{Narg}} + x_{\text{NtrR}}$ |
| S | $x_{\text{CH2}} + x_{\text{OH}}$ |
| T | $x_{\text{CH3}} + x_{\text{CH2}} + x_{\text{OH}}$ |
| V | $2x_{\text{CH3}} + x_{\text{CH1}}$ |
| W | $3x_{\text{CH0}} + x_{\text{CH2}} + 5x_{\text{aroC}} + x_{\text{Ntrp}}$ |
| Y | $2x_{\text{CH0}} + x_{\text{CH2}} + 4x_{\text{aroC}} + x_{\text{OH}}$ |

**Table S3. Per-atom water-to-bilayer transfer energies, $\Delta G_{w,l}^{atom}$**

| Atom Type | $\Delta G_{w,l}$, kcal/mol | $\Delta G_{IMM1}$, kcal/mol | Atom Type | $\Delta G_{w,l}$, kcal/mol | $\Delta G_{IMM1}$, kcal/mol |
|---|---:|---:|---|---:|---:|
| CH0 | -0.7625 | -0.3 | CH2 | -0.7418 | -0.791 |
| CH1 | -0.6889 | -0.3 | CH3 | -1.362 | -0.3 |
| CNH2 | 0.9292 | 0 | COO | 0.3675 | -0.211 |
| S | 0.0386 | 0.612 | SH1 | -0.2582 | 0.612 |
| aroC | -0.4532 | -0.378 | Nhis | 2.2556 | 4.147 |
| Ntrp | 3.3449 | 3.102 | Nlys | 6.7870 | 9.305 |
| NH2O | 0.9292 | 2.773 | Narg | 1.9274 | 3.311 |
| NtrR | 0.9637 | 3.102 | ONH2 | 0.9292 | 2.773 |
| OOC | 0.7351 | 4.206 | OH | 1.4193 | 3.3 |

**Table S4. Full names of lipid types and abbreviations**

| Lipid Name | Full Lipid Name | Head Group Type |
|---|---|---|
| DLPC | 1,2-dilauroyl-sn-glycero-3-phosphocholine | PC |
| DMPC | 1,2-dimyristoyl-sn-glycero-3-phosphocholine | PC |
| DOPC | 1,2-dioleoyl-sn-glycero-3-phosphocholine | PC |
| DPPC | 1,2-dipalmitoyl-sn-glycero-3-phosphocholine | PC |
| POPC | 1-palmitoyl-2-oleoyl-glycero-3-phosphocholine | PC |
| DLPE | 1,2-dilauroyl-sn-glycero-3-phosphoethanolamine | PE |
| DMPE | 1,2-dimyristoyl-sn-glycero-3-phosphoethanolamine | PE |
| DOPE | 1,2-dioleoyl-sn-glycero-3-phosphoethanolamine | PE |
| DPPE | 1,2-dipalmitoyl-sn-glycero-3-phosphoethanolamine | PE |
| POPE | 1-palmitoyl-2-oleoyl-sn-glycero-3-phosphoethanolamine | PE |
| DLPG | 1,2-dilauroyl-sn-glycero-3-phospho-(1'-rac-glycerol) | PG |
| DMPG | 1,2-dimyristoyl-sn-glycero-3-phospho-(1'-rac-glycerol) | PG |
| DOPG | 1,2-dioleoyl-sn-glycero-3-phospho-(1'-rac-glycerol) | PG |
| DPPG | 1,2-dipalmitoyl-sn-glycero-3-phospho-(1'-rac-glycerol) | PG |
| POPG | 1-palmitoyl-2-oleoyl-sn-glycero-3-phospho-(1'-rac-glycerol) | PG |

**Table S5. Water density parameters fit from all-atom molecular dynamics data**

| Lipid Type | Chain | Thickness ($t$) | Pseudo Thickness ($\tau$) | Steepness ($s$) | Temperature (℃) |
|---|---|---|---|---|---|
| DLPC | 12:0/12:0 | 15.35 | 199.57 | 0.343 | 37 |
| DLPE | 12:0/12:0 | 16.45 | 1538.31 | 0.446 | 37 |
| DLPG | 12:0/12:0 | 14.38 | 457.67 | 0.413 | 37 |
| DMPC | 14:0/14:0 | 17.97 | 390.54 | 0.343 | 37 |
| DMPE | 14:0/14:0 | 19.26 | 7906.16 | 0.446 | 37 |
| DOPC | 18:1/18:1 | 18.64 | 707.00 | 0.343 | 37 |
| DOPE | 18:1/18:1 | 19.61 | 3419.03 | 0.446 | 37 |
| DOPG | 18:1/18:1 | 17.97 | 1233.72 | 0.413 | 37 |
| DPPC | 16:0/16:0 | 20.03 | 925.15 | 0.343 | 37 |
| DPPE | 16:0/16:0 | 22.11 | 13436.10 | 0.446 | 37 |
| DPPG | 16:0/16:0 | 18.92 | 1229.49 | 0.413 | 37 |
| POPC | 16:0/18:1 | 19.10 | 815.94 | 0.343 | 37 |
| POPG | 16:0/18:1 | 18.69 | 2379.13 | 0.413 | 37 |

**Table S6. Weights for membrane energy functions based on `score_12`**

| Energy Term | Description | M07 (? ) | M12 (? ) |
|---|---|---|---|
| `fa_atr` | van der Waals attractive energy | 0.78 | 0.8 |
| `fa_rep` | Repulsive energy | 0.43 | 0.44 |
| `fa_intra_rep` | Intra-residue repulsive energy | 0.004 | 0.004 |
| `fa_pair` | Statistical residue-pair interaction energy | 0.48 | 0.49 |
| `fa_dun` | Knowledge-based rotamer energy | 0.56 | 0.56 |
| `fa_mpsolv` | Lazaridis IMM1 membrane solvation energy | 0.60 | 0.35 |
| `fa_mpenv` | Lazaridis IMM1 membrane environment energy | 0.48 | 0.3 |
| `fa_mpenv_smooth` | Knowledge-based membrane environment energy | 0.48 | 0.5 |
| `ref` | Amino acid reference energy | 1.0 | 1.0 |
| `hbond_lr_bb` | Long-range backbone hydrogen bond energy | 1.16 | 1.17 |
| `hbond_sr_bb` | Short-range backbone hydrogen bond energy | 1.16 | 1.17 |
| `hbond_bb_sc` | Backbone to side-chain hydrogen bond energy | 1.16 | 2.34 |
| `hbond_sc` | Side-chain to side-chain hydrogen bond energy | 1.1 | 2.2 |
| `p_aa_pp` | $\phi, \psi$-dependent amino acid propensity | 0.64 | 0.32 |
| `dslf_ss_dst` | Disulfide bonding energy for S-S distance | 0.5 | 0.5 |
| `dslf_cs_ang` | Disulfide bond energy for CSS angle | 2 | 2 |
| `dslf_ss_dih` | Disulfide bond energy for SS dihedral | 5 | 5 |
| `dslf_ca_dih` | Disulfide bond energy for CC dihedral | 5 | 5 |
| `pro_close` | Proline closure energy | 1.0 | 1.0 |
| `rama` | Rakmachandran energy | 0.2 | 0.2 |
| `omega` | $\omega$ torsion energy | 0.5 | 0.5 |

**Table S7. Weights for membrane energy functions based on ref2015**

| Energy Term | Description | R15 (? ? ) | M19 |
|---|---|---|---|
| fa_atr | van der Waals attractive energy | 1.0 | 1.0 |
| fa_rep | Repulsive energy | 0.55 | 0.55 |
| fa_sol | Lazaridis-Karplus solvation energy | 1.0 | 1.0 |
| fa_intra_sol_xover4 | Intra-residue solvation energy | 1.0 | 1.0 |
| lk_ball_wtd | Orientation-dependent solvation energy | 1.0 | 1.0 |
| fa_intra_rep | Intra-residue repulsive energy | 0.005 | 0.005 |
| fa_elec | Coulomb electrostatics energy | 1.0 | 1.0 |
| pro_close | Proline closure energy | 1.25 | 1.25 |
| fa_dun | Knowledge-based rotamer energy | 0.56 | 0.56 |
| hbond_lr_bb | Long-range backbone hydrogen bond energy | 1.0 | 1.0 |
| hbond_sr_bb | Short-range backbone hydrogen bond energy | 1.0 | 1.0 |
| hbond_bb_sc | Backbone to side chain hydrogen bond energy | 1.0 | 1.0 |
| hbond_sc | Side chain to side chain hydrogen bond energy | 1.0 | 1.0 |
| dslf_fa13 | Disulfide bonding energy | 1.25 | 1.25 |
| rama_prepro | Ramachandran energy | 0.45 | 0.45 |
| omega | $\omega$ torsion energy | 0.5 | 0.5 |
| p_aa_pp | $\phi, \psi$-dependent amino acid propensity | 0.6 | 0.6 |
| fa_dun | Knowledge-based rotamer energy | 0.7 | 0.7 |
| yhh_planarity | Tyrosine $\chi_3$ torsion energy | 0.625 | 0.625 |
| ref | Amino acid reference energy | 1.0 | 1.0 |
| fa_water_to_bilayer | Water-to-bilayer transfer energy | 0.0 | 1.0 |

**Table S8. Lipid composition parameters for $\alpha$-helical peptide energy landscape calculations.**

| Target | PDB Code | Experimental Conditions | Parameters | Ref. |
|---|---|---|---|---|
| Acetylcholine M2 segment | 1a11 | DPC micelles | DLPC, 20℃ | (? ) |
| Influenza A M2 channel | 1mp6 | DMPC vesicles | DMPC, 30℃ | (? ) |
| VPU-forming domain HIV-1 | 1pje | DOPC:DOPG 9:1 | DOPC, 30℃ | (? ) |
| NMDA receptor segment | 2nr1 | DPC micelles | DLPC, 20℃ | (? ) |
| WALP23 | NA | DOPC | DOPC, 30℃ | (? ? ) |

**Table S9. Minimum energy orientations of $\alpha$-helical transmembrane peptides**

| Target | M07 $\theta_{min}$ (°) | M07 $d_{min}$ (Å) | M12 $\theta_{min}$ (°) | M12 $d_{min}$ (Å) | M19 $\theta_{min}$ (°) | M19 $d_{min}$ (Å) |
|---|---|---|---|---|---|---|
| 1a11 | -20 | 273 | 1 | 183 | 4 | 26 |
| 1mp6 | -19 | 76 | 0 | 336 | 2 | 137 |
| 2nr1 | -60 | 213 | 19 | 59 | -7 | 222 |
| 1pje | 8 | 66 | -6 | 199 | -1 | 198 |
| WALP23 | 1 | 2 | 0 | 198 | -2 | 32 |

**Table S10. Experimentally measured and predicted $\Delta\Delta G^{\text{mut}}$ values in PagP**

| Mutation | $\Delta\Delta G^{\text{mut}}_{\text{exp}}$ (kcal/mol) (? ) | $\Delta\Delta G^{\text{mut}}_{\text{M07}}$ (REU) | $\Delta\Delta G^{\text{mut}}_{\text{M12}}$ (REU) | $\Delta\Delta G^{\text{mut}}_{\text{M19}}$ (REU) |
|---|---|---|---|---|
| A | 0 | 0 | -0.017 | 0 |
| C | -0.72 | 1.49 | 1.30 | 0.53 |
| D | 2.49 | 5.17 | 5.37 | 3.91 |
| E | 1.18 | 4.00 | 4.34 | 0.37 |
| F | -2.44 | -0.89 | -1.91 | -4.03 |
| G | 1.64 | 2.14 | 2.32 | 4.00 |
| H | 3.32 | 3.38 | 2.34 | 3.44 |
| I | -2.17 | -1.80 | -1.92 | -1.97 |
| K | 3.54 | 3.07 | 2.03 | 4.65 |
| L | -2.01 | -0.50 | -1.41 | 0.91 |
| M | -1.15 | -1.54 | -1.78 | -0.89 |
| N | -2.95 | 2.94 | 3.40 | 3.00 |
| P | 3.82 | 216.14 | 215.09 | 168.83 |
| Q | 2.54 | 1.76 | 1.49 | 1.73 |
| R | 3.22 | 2.73 | 0.45 | 4.02 |
| S | 1.83 | 1.35 | 0.92 | 0.96 |
| T | 0.95 | 0.14 | -0.25 | -1.34 |
| V | -1.75 | -1.95 | -1.00 | -4.20 |
| W | -2.21 | -0.59 | -2.39 | -2.39 |
| Y | -1.02 | 0.36 | -0.56 | -4.35 |

**Table S11. Residuals for predicted $\Delta\Delta G^{\text{mut}}$ values in OmpLA**

| Mutation | M07 (REU) | M12 (REU) | M19 (REU) |
|---|---|---|---|
| A | 0.00 | 0.01 | 0.00 |
| C | 1.57 | 1.43 | 0.88 |
| D | 1.90 | 2.04 | 1.00 |
| E | 2.00 | 2.23 | 0.58 |
| F | 1.10 | 0.37 | 1.12 |
| G | 0.35 | 0.48 | 1.67 |
| H | 0.04 | 0.69 | 0.09 |
| I | 0.26 | 0.18 | 0.14 |
| K | 0.33 | 1.07 | 0.79 |
| L | 1.07 | 0.43 | 2.07 |
| M | 0.28 | 0.46 | 0.18 |
| N | 4.09 | 3.86 | 4.21 |
| P | – | – | – |
| Q | 0.55 | 0.75 | 0.58 |
| R | 0.35 | 1.96 | 0.56 |
| S | 0.34 | 0.65 | 0.62 |
| T | 0.58 | 0.85 | 1.62 |
| V | 0.14 | 0.53 | 1.73 |
| W | 1.15 | 0.13 | 0.13 |
| Y | 0.98 | 0.33 | 2.36 |

**Table S12. Contributions of individual energies to $\Delta\Delta G^{\text{mut}}$ values in PagP predicted by M07**

| mutation | fa_atr | fa_rep | pro_close | fa_pair | hbond_sc | fa_mpsolv | rama | omega | fa_dun | p_aa_pp | ref | fa_mpenv |
|---|---|---|---|---|---|---|---|---|---|---|---|---|
| A104A | 0 | 0 | 0 | 0 | 0 | 0 | 0 | 0 | 0 | 0 | 0 | 0 |
| A104C | 0.835 | 0.786 | 0.481 | 0 | 0 | 0.012 | -0.232 | -0.025 | -0.348 | -0.657 | 0.65 | 0.003 |
| A104D | 0.508 | 0.409 | 0.481 | 0.041 | 0 | 0.526 | 0.001 | -0.025 | 1.366 | 0.126 | -1.72 | 3.47 |
| A104E | -2.23 | 0.212 | 0 | 0 | 0 | 0.537 | -0.219 | 0 | 2.912 | -0.51 | -1.21 | 4.523 |
| A104F | -6.147 | 1.166 | 0 | 0 | 0 | 0.999 | -0.255 | 0 | 3.228 | -0.016 | 0.23 | -0.099 |
| A104G | 2.378 | -0.011 | 0 | 0 | -0.03 | -0.397 | 0.195 | -0.054 | -0.402 | 0.863 | -0.27 | -0.131 |
| A104H | -5.556 | 0.733 | 0 | 0 | 0 | 0.981 | -0.283 | 0 | 3.547 | -0.108 | 0.16 | 3.916 |
| A104I | 0.152 | 1.51 | 0.481 | 0 | 0 | 0.033 | -0.252 | -0.04 | -0.209 | -0.963 | -0.81 | -1.71 |
| A104K | -1.679 | 1.123 | 0 | 0 | 0 | 0.225 | -0.21 | 0 | 0.916 | -0.08 | -0.81 | 3.582 |
| A104L | -2.316 | 1.324 | 0 | 0 | 0 | 0.315 | -0.076 | 0 | 1.682 | -0.008 | -0.26 | -1.172 |
| A104M | -2.286 | 0.181 | 0 | 0 | 0 | 0.305 | -0.3 | 0 | 1.59 | -0.525 | -0.74 | 0.245 |
| A104N | -4.625 | 0.566 | 0 | 0 | 0 | 0.749 | -0.089 | 0 | 3.277 | 0.666 | -1.29 | 3.595 |
| A104P | 0.38 | 41.743 | 84.49 | 0 | 0 | 0.2 | 1.062 | -0.012 | -0.658 | 7.184 | -1.05 | -1.808 |
| A104Q | -2.794 | 0.243 | 0 | 0 | 0 | 0.319 | -0.336 | 0 | 2.939 | -0.484 | -1.37 | 3.256 |
| A104R | -4.325 | 1.121 | 0 | 0 | 0 | 0.867 | -0.374 | 0 | 2.767 | -0.656 | -1.38 | 4.704 |
| A104S | -0.792 | 0.425 | 0 | 0.006 | 0 | 0.259 | -0.16 | 0 | 0.405 | -0.079 | -0.53 | 1.816 |
| A104T | 1.034 | 0.537 | 0.481 | 0 | 0 | -0.04 | -0.267 | -0.025 | -0.4 | -0.753 | -1.32 | 0.898 |
| A104V | 0.461 | 1.19 | 0.481 | 0 | 0 | 0 | -0.321 | -0.04 | -0.437 | -1.187 | -0.76 | -1.332 |
| A104W | -7.846 | 1.07 | 0 | 0 | 0 | 1.326 | -0.159 | 0 | 3.248 | 0.136 | 0.51 | 1.125 |
| A104Y | -6.411 | 1.242 | 0 | 0 | 0 | 1.058 | -0.253 | 0 | 3.156 | -0.064 | 0.11 | 1.521 |

**Table S13. Contributions of individual energies to $\Delta\Delta G^{mut}$ values in PagP predicted by M12**

| mutation | fa_atr | fa_rep | pro_close | hbond_bb_sc | fa_mpsolv | rama | omega | fa_dun | p_aa_pp | ref | fa_mpenv | fa_smooth |
|---|---|---|---|---|---|---|---|---|---|---|---|---|
| A104A | 0.858 | -0.059 | 0 | -1.654 | -0.03 | -0.2 | 0 | -1.142 | -0.201 | -0.17 | 1.636 | 0.965 |
| A104C | -1.2 | 0.12 | 0 | 0 | 0.106 | -0.17 | 0 | 0.57 | -0.022 | 1.54 | 0.353 | 0.005 |
| A104D | -4.123 | 0.277 | 0 | 0 | 0.496 | 0.112 | 0 | 3.817 | 0.41 | -0.1 | 2.844 | 1.639 |
| A104E | -3.601 | 0.606 | 0 | 0 | 0.541 | 0.027 | -0.103 | 3.283 | 0.105 | -0.63 | 2.342 | 1.763 |
| A104F | -6.306 | 1.194 | 0 | 0 | 0.582 | -0.254 | 0 | 3.228 | -0.008 | 0.23 | -0.061 | -0.515 |
| A104G | 2.024 | -0.069 | 0 | -1.654 | -0.143 | 0.125 | 0.064 | -1.142 | 0.311 | -0.5 | 2.042 | 1.34 |
| A104H | -7.566 | 2.013 | 0 | 0 | 0.761 | -0.457 | 0 | 3.381 | -0.281 | 0.89 | 2.698 | 0.922 |
| A104I | -5.14 | 0.748 | 0 | -1.654 | 0.497 | -0.342 | -0.015 | 2.729 | -0.335 | 0.64 | 1.13 | -0.17 |
| A104K | -1.722 | 1.149 | 0 | 0 | 0.13 | -0.21 | 0 | 0.916 | -0.041 | -0.81 | 2.239 | 0.37 |
| A104L | -2.375 | 1.355 | 0 | 0 | 0.183 | -0.075 | 0 | 1.682 | -0.005 | -0.26 | -0.732 | -1.194 |
| A104M | -1.65 | 0.213 | 0 | -1.654 | 0.166 | -0.372 | 0 | 0.719 | -0.277 | -0.67 | 1.344 | 0.394 |
| A104N | -6.469 | 1.803 | 0 | 0 | 0.568 | -0.262 | 0 | 3.111 | 0.107 | -0.56 | 2.498 | 1.73 |
| A104P | -1.258 | 42.63 | 84.49 | -1.654 | 0.318 | 0.947 | 0.106 | -1.141 | 3.433 | -0.31 | 0.805 | 1.451 |
| A104Q | -2.679 | 0.662 | 0 | 0 | 0.208 | -0.207 | 0 | 2.413 | -0.056 | -1.13 | 1.598 | 0.669 |
| A104R | -2.595 | 0.403 | 0 | -1.654 | 0.327 | -0.446 | 0 | 0.954 | -0.343 | -1.31 | 4.131 | 0.921 |
| A104S | -0.812 | 0.435 | 0 | 0 | 0.15 | -0.16 | 0 | 0.406 | -0.04 | -0.53 | 1.135 | 0.325 |
| A104T | -1.372 | 0.528 | 0 | -1.654 | 0.217 | -0.405 | 0 | -0.273 | -0.271 | -0.6 | 2.508 | 1.076 |
| A104V | 0.871 | 0.719 | 0.481 | -1.606 | -0.012 | -0.521 | -0.04 | -1.073 | -0.795 | -0.93 | 0.803 | 1.123 |
| A104W | -9.8 | 2.383 | 0 | 0 | 0.932 | -0.333 | 0 | 3.082 | -0.159 | 1.24 | 0.954 | -0.684 |
| A104Y | -6.577 | 1.271 | 0 | 0 | 0.616 | -0.252 | 0 | 3.156 | -0.033 | 0.11 | 0.951 | 0.197 |

**Table S14. Contributions of individual energies to $\Delta\Delta G^{\text{mut}}$ values in PagP predicted by M19**

| mutation | fa_atr | fa_rep | fa_sol | lk_ball_wtd | fa_intra_rep | fa_elec | hbond_bb_sc | fa_dun | p_aa_pp | ref | fa_wtbe | rama_prepro |
|---|---|---|---|---|---|---|---|---|---|---|---|---|
| A104A | 0 | 0 | 0 | 0 | 0 | 0 | 0 | 0 | 0 | 0 | 0 | 0 |
| A104C | -3.158 | 0.682 | 1.171 | 0.026 | -0.04 | -0.29 | 0 | 0.396 | -0.041 | 1.93 | -0.168 | 0.017 |
| A104D | -1.868 | 0.081 | 4.361 | 0.267 | -0.097 | -0.279 | 0 | 2.134 | 0.694 | -3.471 | 1.762 | 0.319 |
| A104E | -2.108 | 0.303 | 2.955 | 0.261 | -0.372 | -0.461 | 0 | 3.036 | -0.129 | -4.049 | 1.2 | -0.276 |
| A104F | -3.713 | 0.793 | 0.903 | 0.259 | -0.198 | -0.001 | 0 | 1.446 | -0.187 | -0.107 | -2.794 | -0.449 |
| A104G | 1.519 | -0.774 | 0.788 | 0 | 0.035 | 0.058 | 0 | 0 | 0.961 | -0.527 | 0.466 | 1.425 |
| A104H | -3.134 | 0.207 | 2.802 | 0.439 | -0.305 | -0.014 | 0 | 1.845 | -0.274 | -1.625 | 4.001 | -0.505 |
| A104I | -6.498 | 0.835 | 1.104 | 0.493 | -0.317 | 0.187 | 0 | 0.739 | -0.253 | 2.022 | -0.274 | -0.016 |
| A104K | 0.308 | -0.7 | 2.853 | -0.41 | -0.032 | -0.847 | -0.871 | -0.075 | -0.029 | -2.106 | 6.804 | -0.239 |
| A104L | -0.535 | 0.408 | 2.178 | -0.45 | -0.167 | -0.704 | -0.871 | 0.882 | 0.039 | 0.27 | -0.063 | -0.081 |
| A104M | -4.446 | 1.712 | 2.108 | 0.058 | -0.08 | -0.248 | 0 | 1.372 | -0.143 | 0.333 | -1.415 | -0.153 |
| A104N | -2.461 | 0.052 | 3.259 | 0.275 | -0.127 | -0.031 | 0 | 1.507 | 0.453 | -2.665 | 2.625 | 0.111 |
| A104P | 0.635 | 51.563 | -0.86 | -0.487 | 0.139 | 1.363 | 0 | -2.142 | 6.887 | -2.862 | 1.323 | 7.594 |
| A104Q | -2.416 | 0.378 | 2.313 | 0.196 | -0.407 | -0.043 | 0 | 2.822 | -0.104 | -2.776 | 2.122 | -0.365 |
| A104R | 0.34 | -0.712 | 2.847 | -0.329 | -0.036 | -0.868 | -0.871 | 0.626 | -0.219 | -1.486 | 5.169 | -0.444 |
| A104S | -0.672 | 0.324 | 2.785 | 0.066 | -0.368 | -0.49 | 0 | 0.403 | -0.045 | -0.979 | 0.223 | -0.294 |
| A104T | -2.813 | 0.341 | 1.637 | 0.049 | -0.413 | -0.406 | 0 | 0.33 | -0.13 | -0.173 | 0.53 | -0.298 |
| A104V | -3.652 | 0.586 | -0.059 | 0.037 | -0.169 | -0.016 | 0 | 0.284 | -0.537 | 1.318 | -1.485 | -0.51 |
| A104W | -5.558 | 0.454 | 1.451 | 0.285 | -0.39 | -0.038 | 0 | 1.471 | -0.045 | 0.936 | -0.942 | -0.034 |
| A104Y | -3.656 | 0.789 | 0.721 | 0.279 | -0.228 | 0.032 | 0 | 1.251 | -0.202 | -0.107 | -2.786 | -0.468 |
